# Preferential generation of pathological tau species in specific subtypes of entorhinal neurons: implications for Alzheimer’s Disease

**DOI:** 10.64898/2026.02.05.703778

**Authors:** I Martinsson, V. Di Maria, M. Carvalho, A. Kobro-Flatmoen, M.L. Potenza, M.P. Witter, C. Kentros

**Affiliations:** Norwegian University of Science and Technology (NTNU)

## Abstract

Alzheimer’s Disease (AD) is distinguished by the presence of two key pathological features: amyloid plaques, accumulations of proteolytic products of Amyloid Precursor Protein, and neurofibrillary tangles (NFTs), intracellular aggregations of microtubule-associated protein tau. NFTs first appear in particular entorhinal cortex (EC) neurons (called pre-alpha neurons) in asymptomatic patients, then continue to spread through other connected brain regions as the disease progresses. This stereotypical progression of tauopathy through synaptically connected brain regions (i.e. Braak stages) not only suggests that the tauopathy in AD spreads transsynaptically, it also raises the question whether particular neuronal subtypes and/or brain regions are especially prone to tauopathy. We explored this question by overexpressing wildtype human tau protein (hTau) in a variety of entorhinal and neocortical neuronal subtypes. We then compared the tendencies of different neuronal cell types to develop different pathological tau species over time and found that tau pathology does indeed develop at markedly different rates in different neuronal subtypes. Perhaps unsurprisingly, the EC is particularly prone, with the likely rat cognates of the pre-alpha neurons (ECLII fan cells) among the first to express pathological tau label. Fan cells were not, however, the neurons with the greatest vulnerability to generate pathological tau species: subsets of ECLIII neurons were found to express disproportional amounts of pathological tau. This is of particular interest given that the next structures to develop AD-related tauopathy after the EC in patients are CA1 hippocampus and subiculum, where ECLIII neurons project, rather than the dentate gyrus and CA3, where ECLII fan cells project. These results demonstrate differential susceptibility of different neuronal subtypes to pathological tau species and suggest distinct roles for different entorhinal neuronal subtypes in the propagation of the tauopathy underlying AD-related neurodegeneration.

## Introduction

The two most prevalent neurological disorders in the Western world, Alzheimer’s Disease (AD) and Parkinson’s Disease (PD), are both incurable progressive neurodegenerative diseases, but their clinical presentation is very different. Neurodegeneration basically means neurons die, so arguably the most important difference between these two diseases is *which* neurons die. While in PD the affected celltype is well known (dopaminergic neurons of the substantia nigra pars compacta), AD’s dual pathological signature of amyloid plaques (extracellular aggregations of proteolytic peptides derived from Amyloid Precursor Protein (O’Brien and Wong 2011)) and neurofibrillary tangles (NFTs, intracellular aggregations of hyperphosphorylated Microtubule-Associated Protein Tau (Grundke-Iqbal et al. 1986)) complicates the matter. Both pathological signs have a stereotypical anatomical progression, but only NFTs seem to correlate closely with cognitive decline (Guillozet et al. 2003; Ossenkoppele et al. 2019). Unlike plaques, which initially form in the frontal-parietal neocortex (Thal et al. 2002); pathological tau species, NFTs and the ensuing cell death are first seen in prodromal patients in the projection neurons of layer II of the transentorhinal cortex (Braak Stage I), which they called pre-alpha neurons (Braak and Braak 1991). The tauopathy then progresses through the rest of the entorhinal cortex (EC; Stage II) and the hippocampus proper (HPC, Braak Stage III) as cognitive decline increases, only reaching the neocortex in later stages. Both the HPC and its deeply interconnected primary input, the EC, are strongly implicated in learning and memory, explaining why AD’s first symptoms tend to be memory-related.

The fact that the tauopathy in AD spreads between synaptically connected brain regions and that it can be “seeded” by injecting patient tissue in animal brains (Clavaguera et al. 2013) led to the “prion” hypothesis in which AD tauopathy spreads trans-synaptically via a prion-like misfolded tau (Clavaguera et al. 2009). The transynaptic spread of the tauopathy in AD is now generally accepted in the field (Walsh and Selkoe 2016; Lewis and Dickson 2016), but significant questions remain, not least why it almost always starts in what the Braaks called pre-alpha cells of the human transentorhinal cortex. The human brain remains largely experimentally inaccessible at the cellular level, so one must look for homologous structures in other animals to explore these questions (Schön et al. 2022). While this remains an “inexact” science, luckily the entorhinal cortex is relatively old cortex which appears to be quite evolutionarily conserved. Given this, the best cognate in non-human primates- and, indeed, in rodents-would be the projection neurons of Layer II of the Lateral Entorhinal Cortex (LEC LII), called “fan cells” due to their fan-like arborization. For instance, LEC LII fan cells are one of the few excitatory neurons that continue to express Reelin in the adult brain (Kobro-Flatmoen, Nagelhus, and Witter 2016), a fact made particularly interesting by the recent finding of a protective allele in the gene (Lopera et al. 2023). However, while the EC and HPC are densely interconnected, the location of the first NFTs found outside the EC in patients is CA1 and subiculum, but LEC LII fan cells project to CA3 and DG. These observations suggest that the anatomically stereotypical progression of NFTs may have a cell-autonomous aspect. In this view, different neuronal cell types may simply have different timescales of vulnerability towards AD tauopathy, giving the appearance of a transsynaptic progression (Walsh and Selkoe 2016). Regardless of which mechanism is more correct (and they are not mutually exclusive), it is reasonable to assume that a neuronal subtypes’ tendencies towards the generation of pathological tau species is affected by the cellular microenvironment (Anand et al. 2025; Roussarie et al. 2020; Fu et al. 2019).

This led us to investigate whether different neuronal cell types are differentially susceptible to the generation of pathological tau species induced by overexpression of wildtype human tau protein (hTau) in a wide variety of neuronal celltypes in the dorsal LEC and surrounding neocortical areas. Since hTau overexpression would eventually lead to pathological tau phosphorylation and aggregation in any cell type, quantifying expression levels in different celltypes of interest over time is critical. We therefore injected a vector designed to express both GFP and hTau in all neuronal types (hSyn 4RhTau-2A-GFP AAVs, see Methods) near the rhinal fissure of wildtype rats such that all neuronal cell types of LEC and surrounding neocortex (mostly the Perirhinal cortex, BA 35 and 36) would be infected. Since the 2A element yields 1:1 stoichiometry between the two proteins (Trichas, Begbie, and Srinivas 2008), one can use GFP intensity as a proxy for hTau expression levels. We thereby compared the time course and development of pathological tau species versus GFP levels and found that different neuronal subtypes do indeed have markedly different susceptibility to the generation of pathological tau species.

## Results

### Experimental Design

To address the question of differential vulnerability of neuronal subtypes to AD-related tauopathy, one needs to model it similarly and quantifiably across a variety of neuronal types. Though there are a large variety of transgenic AD rodent models based upon adding one or more human familial Alzheimer’s (fAD) alleles, none recapitulates every aspect of disease progression. This is especially true for the development of neurofibrillary tangles (NFTs) composed of pathological tau species, let alone its stereotypical anatomical progression seen in patients: no rodent model purely based on familial APP/PSEN mutations develops NFTs. This has led several groups to express mutant human tau species (Lewis et al. 2000; Allen et al. 2002; Pooler et al. 2013), but these arguably model the diseases they underlie (e.g. Fronto-temporal dementia,) better than AD, which involves aggregations of *wildtype* human tau, and has little or no linkage to the MAPT gene apart from rare cases (Strang, Golde, and Giasson 2019; Coppola et al. 2012). Other groups have successfully obtained AD-related pathological species by introducing human tau either from patient extracts or artificially fibrillized Tau (tau “seeding”(Clavaguera et al. 2013; Langer Horvat et al. 2023a)) or via expression with viral vectors (Tetlow et al. 2023; Dujardin et al. 2014). While there are pros and cons to each, we chose a viral approach because it allowed us to explore the response of a variety of neuronal cell types to the quantifiable overexpression of wildtype human tau protein in wildtype (Wistar) rat brain. We focused this “tau challenge” upon the entorhinal cortex as it is the first region to develop NFTs in patients and used co-expression of GFP to aid quantification. The null hypothesis would be a roughly linear relationship between the level of human tau expression and the development of pathological tau species over time regardless of cell type: the more hTau, the more pathological tau species. Any deviations from this would mean that the neuron type has a significant influence on the generation of pathological tau species.

### AAV Injections induce similar hTau and eGFP expression in rhinal cortices

Figure 1 shows the basic experimental protocol employed to determine whether different neuronal cell types vary in their predilection to generate pathological tau species (see methods section for details). Since tau overexpression can lead to pathological tau species in many cell types (Kahlson and Colodner 2015), we generated AAVs that result in 1:1 stoichiometry between hTau and a much more easily measured fluorophore, aiding the quantification of hTau expression levels. Large volumes of these hTau-AAVs were injected near the rhinal fissure of month-old wildtype Wistar rats such that the various neuronal subtypes of both lateral entorhinal cortex (LEC) and adjoining neocortical areas would be infected (see Methods for details). Rats were then sacrificed at 1, 2 and 5 months post injection (mpi) and brains sectioned for histology with antibodies raised against pathological tau species **(Fig 1a and b**). Scans of the resulting slides were anatomically segmented and the amount of signal per cell was determined for each antibody for subsequent quantitative analysis (see Methods). As can be seen in Figure 1c, such injections lead to widespread GFP expression in many cell types of the rhinal cortices, both the Lateral Entorhinal Cortex, and overlying neocortical regions. As each injection is unique, we examined the extent of spread of GFP expression in the LEC and adjacent cortices. To be included in our analysis the injection had to include each of our regions of interest, i.e. neurons in Layers I-VI of the LEC as well as adjoining neocortical areas (**Fig 1c**, **supplemental fig 1**), though other regions were sometimes included (See **supplemental Table 1** for description of injections). As expected, the neurons that express GFP also express 4R-hTau (**Fig 1c, d, and e**). GFP expression can be seen throughout the LEC and adjacent cortices (**Fig1c-e**), and injections into S1 also yield GFP positive neurons with HT7 expression (**Fig 1f**). LEC LII fan cells are clearly expressing more GFP in most of our samples (**Fig1c, Supplemental fig 1**). This is likely due to an apparent preference at the infection level, presumably due to interactions with the AAV capsid. This underscores the need to use GFP to normalize for visualizing celltype-specific effects: the key question is what percentage of hTau becomes pathological tau over time.

**Figure 1.**
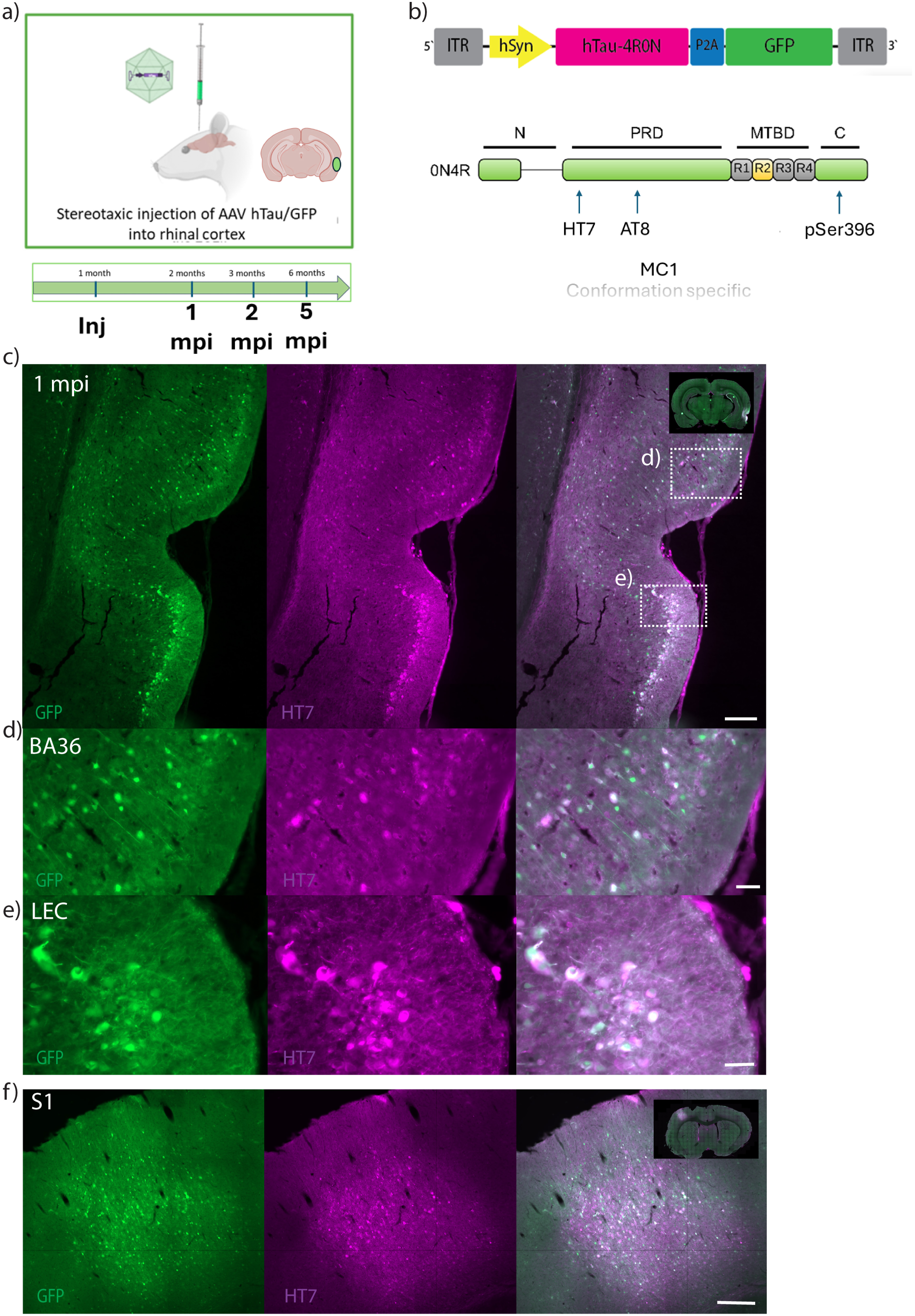
Local injections of hTau-2a-GFP-AAV below the rhinal fissure lead to stochiometric expression of human Tau and GFP. **a)** schematic of experimental design. 30 days old Wistar rats injected just below the rhinal fissure then sacrificed at 1MPI, 2 MPI or 5 MPI for IHC. **b)** Schematic of viral construct (top) and of 4R-hTau protein with annotated antigen recognition sites for antibodies used in this study.**c)** Representative micrograph of injection near the rhinal fissure in 1mpi animal. Inset shows whole-hemisphere scan from the same section. scalebar = 200um, note, hTau expression above and below the rhinal fissure **d)** Enlarged inset from **c)** showing HT7 expression in GFP positive neurons in the ectorhinal cortex (BA36) scalebar 50um **e)**, Enlarged inset from c) showing HT7 expression in GFP positive neurons in LEC, scalebar 50 um. **f)** Micrograph shows injection into S1 (somatosensory cortex), Inset shows whole-hemisphere scan from same section. Scalebar = 100um

#### Region and celltype-specific development of pathological hTau species

One sees a very different picture when comparing GFP signal to that of antibodies against pathological tau species (**Fig2a**). As can be seen in Figure 2, one starts to see label with AT8, the phospho-tau antibody commonly used to diagnose AD, after only a single month post-injection. Whereas the GFP (and thereby hTau) is spread throughout the rhinal cortices, the vast majority of AT8 label is restricted to the superficial LEC (**Fig 2c, Supplemental fig 2**), especially to the Reelin-expressing LEC LII fan cells, though there is also scattered positive staining in LEC LIII (**Fig2d**). GFP intensity is a good predictor of AT8+ as expected, but this relationship varies significantly by neuronal cell type. For instance, many neocortical neurons express GFP intensely but are basically devoid of AT8 label (**Fig2b, Supplemental Fig 2**). At 2mpi the picture looks similar in the LEC, but AT8 also starts appearing in BA35 in neurites and cell bodies (**Fig 2d**). At 5 months post-injection, curiously, there appears to be less cell body signal in LEC LII, but much more AT8 signal throughout neurites and neurons across the rhinal fissure (**Fig 2d-e**, **Supplemental fig 3**), and the Reelin labeling appears less robust in the LEC of 5 mpi animals (**Fig2d**). At 1mpi little or no AT8 signal is seen in neocortical neurons despite robust GFP (and hTau) signal, both in overlying rhinal cortices and for control animals injected in the primary somatosensory cortex S1 (**Fig 1b, f, Supplemental Fig 2** (i.e. ectorhinal (BA36) and S1). Consistent with their late appearance in AD, S1 neurons remained largely unaffected by at 2 and 5mpi (**Supplemental Fig 2**).

**Figure 2.**
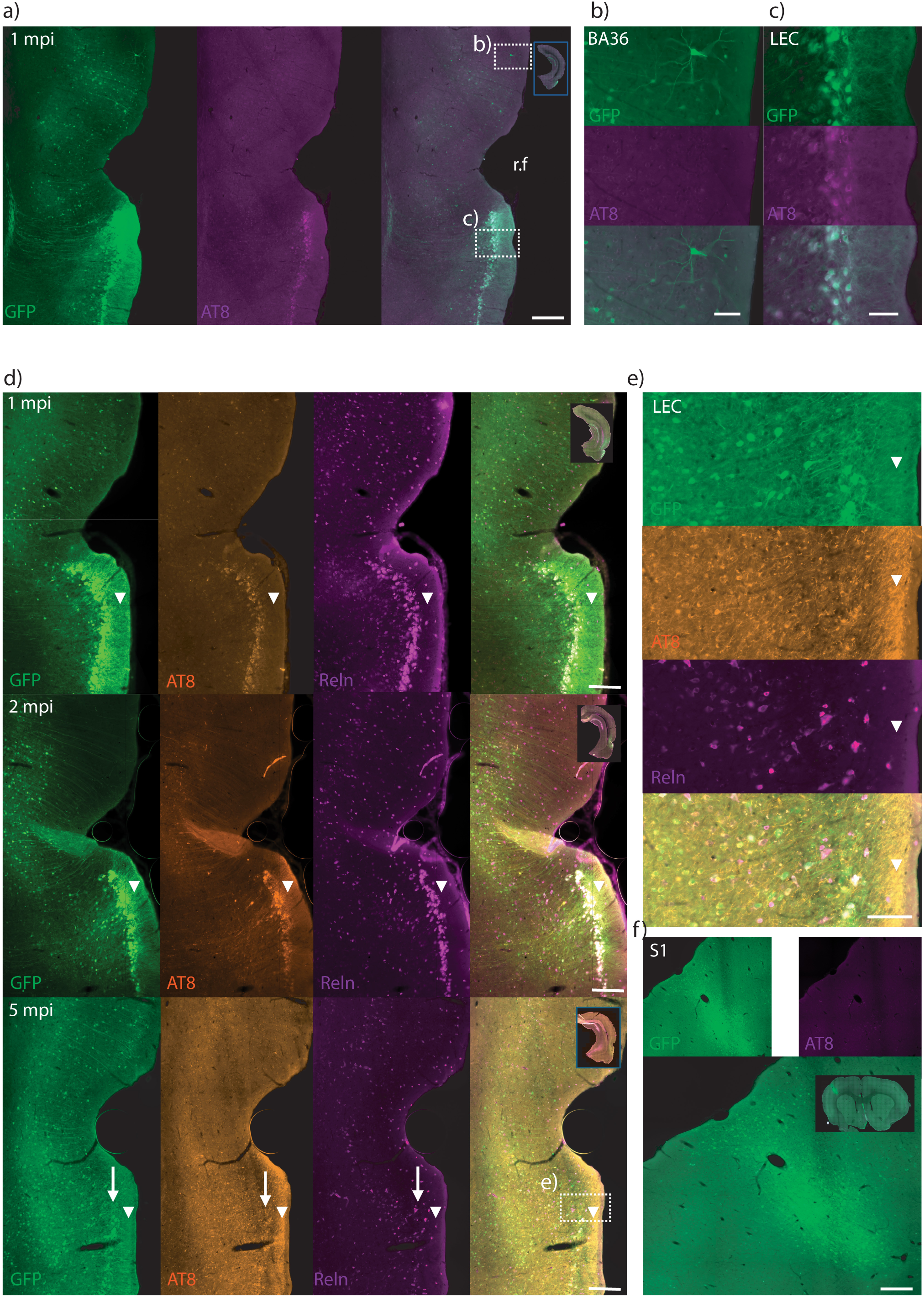
Early phospho-Tau antibody AT8 is concentrated in LEC, primarily in LEC LII fan cells. **a)** Representative micrograph of 1 MPI animal injected near the rhinal fissure. Inset shows whole-hemisphere scan, scalebar= 100um **b)** enlarged inset from **a)** showing that AT8 phospho-tau labeling is largely absent in perirhinal cortex cortex (BA36) despite there being GFP positive neurons. Scalebar = 50 um. **c)** inset from A) showing layer specific labeling of AT8 primarily in layer II of LEC. Scalebar = 50 um. **d)** micrographs from 1, 2 and 5 mpi animals labeled with AT8 and reelin. Note progression of AT8 staining from primarily reelin positive neurons at 1 MPI to LEC LIII and individual perirhinal neurons at 2 MPI to more extensive labeling of processes as well at 5 MPI. Also note an apparent reduction in LEC LII reelin signal at this timepoint highlighted by the arrow. Arrow heads highlight LEC LI, note the lack of AT8 at 1 and 2 mpi compared to 5 mpi. Scalebar = 200 um. **e)** enlarged inset from **d)** The arrow points to AT8 labeling in neurites and parenchyma of layer III. The arrowhead points to intense AT8 labeling in processes in layer I. Scalebar = 100um **f)** Micrograph of S1 injection labeled with AT8, inset shows whole-section scan. Despite robust GFP signal, AT8 is largely absent. Scalebar = 100um

#### Different neuronal subtypes are differentially vulnerable to distinct pathological tau species

The development of AD-related tauopathy is a complex but well-studied process with literally dozens of antibodies raised against different AD-related antigens ranging from patient brain extracts to phosphopeptides to particular oligomeric and conformational states(Petry et al. 2014). Each are thought to model different stages or aspects of the development of the tauopathy in AD, raising the question of whether the tauopathy happens the same way in different cells, or whether different neuronal subtypes differ in their susceptibility to develop these different pathological tau species. We explored this by immunohistochemistry with antibodies against three different pathological tau antigens (see Figure 1b): AT8, an early phospho-tau antigen in the Proline-Rich domain of tau; MC1, an antibody specific to a particular pathological conformation of tau(Jicha et al. 1997), and pSer396, a phospho-antigen on the C-terminal domain of tau thought to appear later in the pathological cascade towards NFTs (Su, Cummings, and Cotman 1994)). As seen in **Figure 3**, each of these distinct pathological tau species tend to develop primarily in LII and LIII of LEC, where one sees the only appreciable signal at 1MPI (**Fig3**). At 1 MPI most GFP positive neurons in ECLII are also AT8 positive, but MC1 reactivity is much sparser, showing up in just a few GFP+ neurons in LEC LII and LIII (**Fig3a**). At 2MPI entorhinal label still dominates, but one can see label in other cell types as well, with both AT8 and MC1 present in the deeper layers of BA35 (**Fig 3a**). At 5 MPI the signal of both AT8 and MC1 shifts mostly to the processes of neurons (**Fig 3c**), which complicates the detection of cell bodies for quantification. Consistent with its later appearance in AD-related tauopathy, pSer396 is barely detectable at 1MPI, although some neurons in LII LEC do label weakly (**Supplemental figure 4a-b**). However, at 2 MPI still quite faint pSer396 label is found exclusively in neurons of LEC LII and LIII (**Fig3d and e**). When one compares the intensity of these antigens to GFP it becomes clear that there is a subpopulation of LEC LIII neurons with relatively weak GFP label but very strong label with these pathological tau species. At 5 MPI, pSer396 staining is prominent in processes around the LEC extending across layers (**Fig 3d and f**), consistent with movement to axons of LECLII and LIII neurons. Thus, although at 5 MPI all antibodies label both LEC LII neurons and LEC LIII, in LEC LIII label for MC1 and pSer396 is at particularly high levels relative to both the GFP and AT8 label (**Fig 3d, e and f**), demonstrating that the tendency towards developing particular pathological tau species is a property of specific neuronal subtypes.

**Figure 3.**
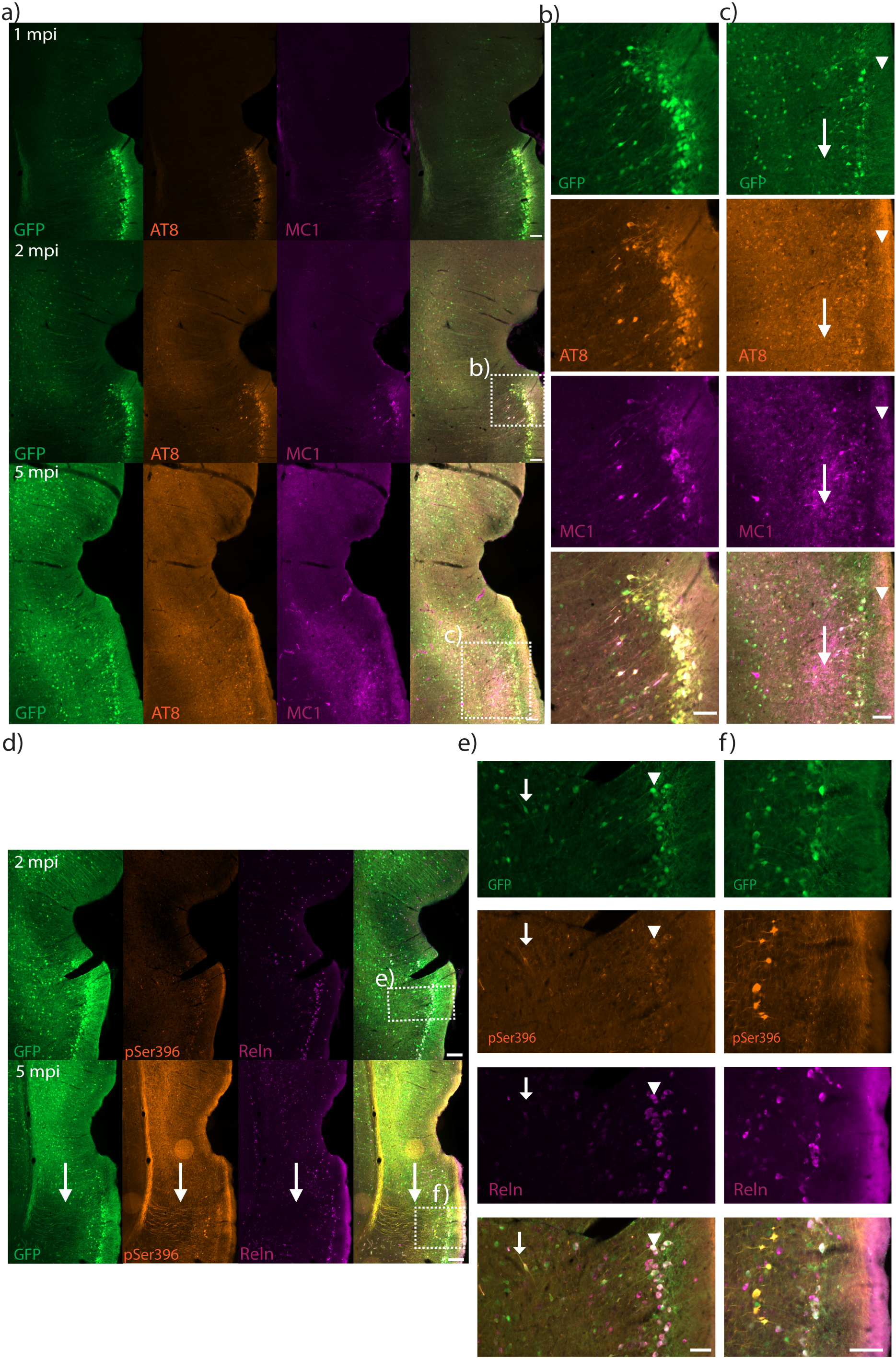
LEC neuronal subtypes are differentially prone to develop distinct pathological tau species. **a)** Representative micrographs of 1,2 and 5 MPI animals labeled for AT8 and MC1. Note that the conformation-specific MC1 antigen is expressed primarily in LIII and LII of LEC at 1 and 2mpi, but at 5mpi both antigens are present in neuropil and show a more diffuse labeling. **b)** Inset from **a)** highlighting LEC LII and LIII AT8 and MC1 positive neurons. Scalebar = 50um. **c)** Enlarged inset from **a)** showing AT8 and MC1 going into the neurites at 5mpi. Arrow points to AT8 and MC1 positive neurites in layer III whereas arrowhead points to layer I neurite labeling, scalebar = 50um. **d)** Micrograph from 2mpi and 5mpi animals labeled with pSer396 and Reln showing that at 2MPI the pSer396 labeling is relatively weak and found mainly in LEC LII, but some LIII neurons are labelled as well. At 5 mpi pSer396 intensity is substantially more widespread, with multiple neurites and areas affected. Arrow points to Temporo-ammonic pathway pSer396 labeling. Scalebar = 200um. Also note the sparse Reln labeling in LII at 5mpi. **e)** inset from **d)** showing labeling in neurons of LEC LII and LIII. Arrow points to Reln expressing neuron in LIII expressing pSer396 at 2mpi whereas arrowhead points to Reln positive neurons in LII that are pSer396 positive. Scalebar = 100um. **f)** lower inset from **d)** showing intense labeling of pSer396 in LEC LII and LIII at 5mpi. Scalebar = 100um

#### Different neuronal subtypes form different pathological tau species at different rates

To quantitate these data, we used scans of HT7, AT8 and MC1 labeled sections from 1 mpi and 2 mpi and used a semi-automated counting algorithm (see Methods) to estimate signal intensity for each GFP+ cell, resulting in 14227 GFP+ neurons from 6 injected rats. The intensity of tau and GFP were normalized by animal and antibody, so the lowest intensity objects were adjusted to 0 and the highest to 10. Note that this leads to a significant floor effect in the GFP channel, as we are only counting cells with appreciable GFP signal. As we saw clear anatomical differences in the predilection to develop pTau species, we manually delineated five anatomical regions (LEC LII, LEC LIII, LEC LIV-VI, BA35 and BA36) in the GFP channel as a first pass at quantifying Tau vulnerability in different neuronal subtypes.

**Figure 4a** shows a summary scatterplot of all detected GFP positive ROIs with colors to highlight the different antibodies used. Trend lines per antibody show that overall HT7 shows a linear 1:1 relationship with hTau, compatible with a X=Y relationship with GFP. The regressions for AT8 (at 1MPI p = 0.046)(2MPI trend p = 0.06) and MC1(0.005 and 0.001 respectively for 1 and 2 MPI) were not consistent with a 1:1 hTau GFP relationship (**Supplemental table 2**). Thus, we can reject the hypothesis that the amount of hTau expression is the most important explanation for the development of pathological tau species. To investigate neuronal subtype sensitivity to hTau challenge, we classified data points based on the residuals to a hypothetical X=Y linear tau relationship, leading to classes of high residuals, low residuals and linear-like data points. These were then further subdivided into Linear low-Tau, Linear mid-Tau, Linear High-Tau, Above High-Tau, Above low-Tau and Below, figure 4b shows the cut-offs for the different classes highlighted in the graph (**Fig 4b**). ROIs were also classified post-hoc by the “parent” region of origin, i.e which anatomical region the object was detected in (**Fig 4c**). The wildtype hTau (HT7) label correlates very well with the GFP label at 1MPI, with a relatively tight distribution and a slope near 45 degrees regardless of anatomical subregion, though at the 2 mpi timepoint the fit between was less clear, possibly because conversion to certain pathological tau species occludes this epitope (Langer Horvat et al. 2023b)(See **Supplemental table 2** for linear regression fits). In contrast, the AT8 and MC1 pathological tau antibody signal significantly deviates from a 1:1 linear relationship in an anatomically-specific manner. Notably, Supralinear High-Tau (Above High-Tau) was present mostly in LEC LII and LEC LIII at 1 mpi, while at 2mpi more regions contribute to this class (**Fig 4d**). This whereas the class we labeled as linear High-Tau is mainly populated by LEC LII at both 1mpi and 2 mpi for AT8 and MC1. At 2MPI the perirhinal cortex (BA35) became more involved, particularly the deeper layers (**Supplemental figure 5**). As pSer396 was not appreciable in most 1MPI brains, we did not quantitate them.

**Figure 4.**
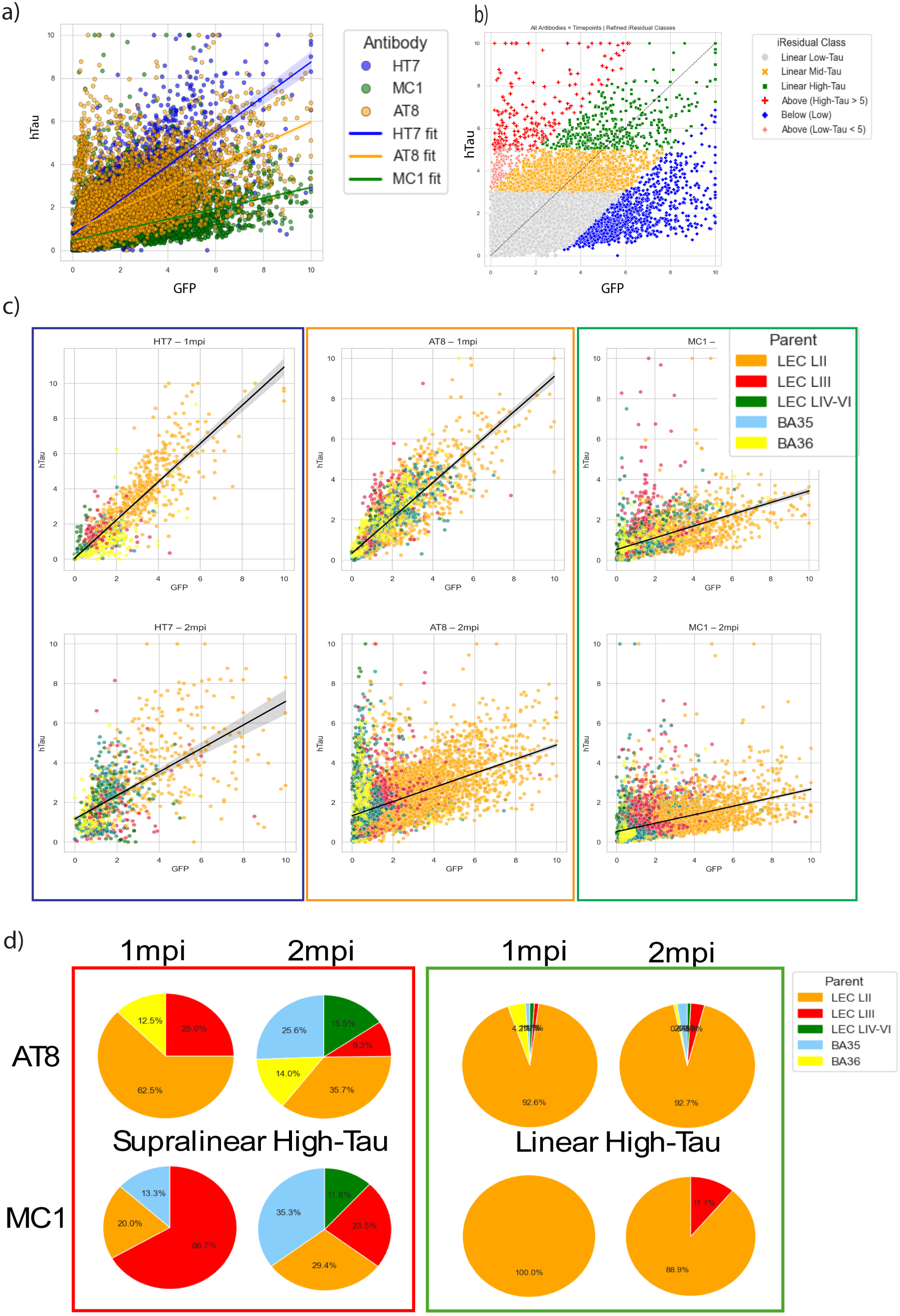
Automated quantification of pathological tau species vs GFP. **a)** Scatterplot of normalized hTau and GFP fluorescence intensity values from QuPath detected “Cell” objects. Wildtype hTau (HT7,Blue), AT8(Orange) and MC1(Green) all cluster differently and linear regression lines from bulk of all data per antibody is highlighted in the figure. Note that all three antibodies have differential slopes with substantial amount of residuals, the data in the scatterplot comes from 3 animals per timepoint and antibody and 1000-8000 cells per antibody/timepoint. **b)** Graph shows scattered data classified based on non-linearity of hTau expression and deviation from a hypothetical 1:1 relationship with GFP. **c)** Subplots from **a)** with objects color coded by “parent” origin i.e. the anatomically segmented region the object was detected in. **d)** Pie charts of the objects sorted into the classes Supralinear High-Tau (Above High-Tau) and Linear High-hTau. At 1mpi both AT8 and MC1 are dominated by LEC LII and LIII for the supralinear hTau objects. Whereas, 2MPI shows contribution from more areas, notably BA35. In contrast, the linear high-hTau class is dominated by LEC LII for all antibodies and timepoints.

Although counts done in this manner are unbiased by the experimenter and can thereby highlight potentially overlooked regions and populations, there are issues with automated counting of immunohistochemical data. One complication is the presence of signal in neurites and/or neuropil. This becomes increasingly prevalent at later stages (5mpi), and can lead to overestimation of pTau presence from neurites overlapping a detected cell body, or underestimation due to high “background” signal. Indeed, we suspect some of the signal in BA35 is due to increased staining of dystrophic neurites in the deeper layers of BA35 at the 2mpi timepoint (**Supplemental figure 5, 6 and 7**)). The other potential complication is epitope loss with disease progression which could explain the loss of 1:1 linearity in HT7/GFP at 2mpi as it has been shown that the HT7 epitope can be occluded by pathological tau species(Langer Horvat et al. 2023a).

Though imperfect, the scatterplots show a number of quite clear trends. Most importantly, the vast majority of neurons that turned their wildtype hTau into appreciable amounts of pathological tau species in 1-2 months were in Layer II of LEC (Orange and Red color **Figure 4c and d**). Surprisingly, LEC LII neurons were not, however, as sensitive to hTau expression in a quantitative sense as neurons in LEC LIII. The connectivity of LEC LIII neurons is of particular interest as they are not only densely interconnected with LEC LII fan cells (Rowland et al, 2011; Butola et al, 2025), they project to CA1 and Subiculum, unlike fan cells, which project to Dentate Gyrus and CA3.

#### Cellular Identity of hTau Hypersensitive LEC Neurons

While the above layer-based analysis shows that neurons in LEC Layers II and III are much more likely to develop pathological tau species than neocortical neurons, even within a single layer of LEC there are several different cell types. Layer II of LEC can be divided into two sublayers: LIIa, composed largely of Reelin-positive fan cells; and LIIb, composed of pyramidal neurons positive for both Calbindin and Wolframin Syndrome 1(WFS1) (Leitner et al. 2016). Due to the curvature of the entorhinal neuraxis it can be difficult to disambiguate LEC layers IIa from IIb, and IIb from III, so we stained representative sections with the above markers (**Fig 5a**). AT8 label is primarily found in the calbindin-negative, Reelin-positive neurons of LIIa, rather than in LIIb. The very strongly AT8-positive neurons below LEC LIIa are calbindin-negative, confirming that they are LIII neurons, not LIIb pyramidal cells. MC1 is detectable in calbindin-negative neurons in LEC LIII and LIIa, but GFP-positive, WFS1-positive neurons in LIIb did not stain with MC1 (**Fig 5b**).

**Figure 5.**
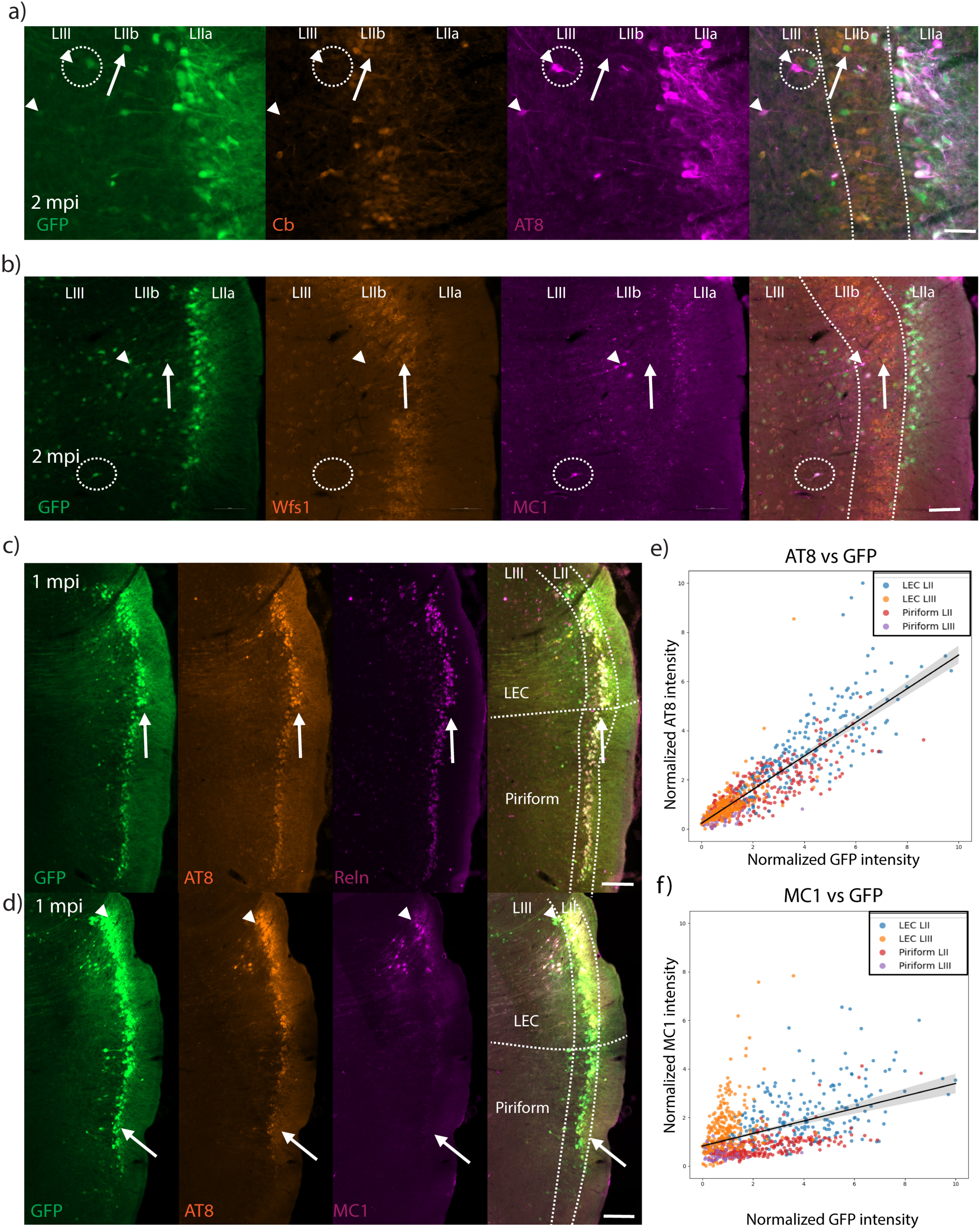
Identification of neuronal cell types hypersensitive to hTau overexpression. **a)** Representative micrograph of 2 MPI animal labeled for AT8 and Calbindin d28k(Cb). Scalebar = 50um. Arrow heads show AT8 positive CB negative neurons in LIII, note that the circled arrow point highlights a neuron with undetectable GFP. The upward-pointing arrow indicates a GFP positive, calbindin-positive neuron in LIIb that is negative for AT8. **b)** Micrograph from 2 MPI animal labeled with WFS1 and MC1. Arrowhead indicates a putative MC1-positive beaded neurite, circled is a strongly labeled MC1 positive neuron in LIII that is WFS1 negative. The upward-pointing arrow points to GFP- and WFS1-positive neuron bordering LIIb without any MC1 labeling. Scalebar 50 um. **c)** Micrograph of a 1 mpi animal labeled with AT8 and reelin from section at the LEC-Piriform boundary, denoted by an arrow pointing at a “notch” in the reelin labeling. Note that reelin positive neurons in LII of LEC and piriform both show AT8 label, and it is even detectable in LIII of both structures (though much more robustly in LEC). Scalebar = 200 um. **d)** Micrographs from a 1 MPI animal labeled with AT8 and MC1. The arrowhead highlights neurons in dorsal LEC, which express high levels of GFP, AT8, and MC1, in contrast the arrow points at Piriform LII neurons which express GFP and AT8, but not MC1. Scalebar = 200 um. **e)** Scatterplot from the 1mpi animal injected sufficiently rostral to obtain GFP label in the piriform cortex. Note that AT8 follows a linear relationship with GFP with a few outliers in LEC LII and LEC LIII. **f)** Similar scatterplot for MC1, note that unlike AT8, MC1 clusters differentially for LEC LII, LEC LIII and Piriform cortex, with the vast majority of label coming from LEC II and III, the very high GFP expression levels in some LIII piriform neurons notwithstanding (Figure 5d).

Multiple lines of evidence converge on a role for Reelin in the development of AD (Kobro-Flatmoen, Nagelhus, and Witter 2016; Kobro-Flatmoen et al. 2023; Lopera et al. 2023), so clearly Reelin is a prime candidate for why these particular neurons develop AD-related tauopathy. It is certainly true that the vast majority of AT8+ neurons are also Reelin-positive (due almost entirely to LEC LII fan cells), but they are also overrepresented in our sample, likely due to the attractiveness of these large, highly glycosylated neurons to the viral capsid. While we aimed for LEC and dorsally adjacent BA 35 and 36 with our injections, in some cases the injections were more rostral, thereby inadvertently performing an hTau challenge in the piriform cortex as well. As the neurons of piriform layer II are one of the few other excitatory neuron types that are Reelin-positive in adults (Carceller et al. 2016), this provided a means to investigate whether other Reelin-positive neurons are equally susceptible to tauopathy. We should note that these Reelin-positive piriform neurons also appear to also have an affinity for the viral capsid, as their GFP signal was higher than most other neurons, though not as high as LEC LII neurons. Interestingly, LII piriform neurons also strongly express AT8 at 1MPI (**Fig 5c**), arguing for a role for Reelin in the development of tauopathy. However, MC1 label was only detectable in LEC neurons in both layers II and III (**Fig 5d**). Thus, even though the Reelin positive neurons of layer II LEC and Piriform cortex both express high levels of GFP and AT8 label, the MC1 label appears to be specific to entorhinal neurons, at least at this early timepoint. We quantified this by counting signal of neurons in LII and LIII of LEC and Piriform cortex in this animal. Interestingly, AT8 more or less showed a linear relation with GFP (**Fig 5e**) in these neurons, whereas MC1 showed a flat line for the most part in Piriform neurons and scattered populations of MC1 positive objects in LII and LIII of LEC (**Fig 5f**). Thus, while Reelin might play a part in the vulnerability of ECLII neurons to AD-related tauopathy, it is not the whole story: there are clearly other relevant features of superficial entorhinal neurons.

#### Pathological tau species in axonal projections

In AD, pathological tau species first appear in dendrites and soma and then migrate to axons (Braak and Del Tredici 2018). We see the same basic progression in our experiments, with label migrating to the EC neurons axons projecting to the hippocampus. Injections into LEC lead to visible GFP positive projections in the hippocampal formation (**Fig6a-d**). At 1 MPI both GFP and HT7 are present in the terminals of LIIa, and LIIb and LIII neurons (**Fig6e**). At this timepoint, pSer396 is not present in the processes (**Fig6f**). Interestingly, at 5 MPI, both HT7 and pSer396 are present in the hippocampal projections (**Fig6g-h**). Note, that the projections to the Stratum lacunosum moleculare (SLM) of CA1 appear particularly strong for pSer396 **(Fig 6i**), and that the labeling in the Molecular layer of the dentate gyrus is strongest in the medial part. While the labeling pattern in SLM does not exclude the contribution of LIIb neurons, it corroborates the presence of late-stage pTau species in LIII neurons. Figure 6i shows inset from 6h highlighting the appearance of pSer396 positive process bundles in SLM CA1. The presence of pSer396 at the terminal fields of the LEC LIII and LIIa output provide further evidence that these two layers are affected early in the pathology.

**Figure 6.**
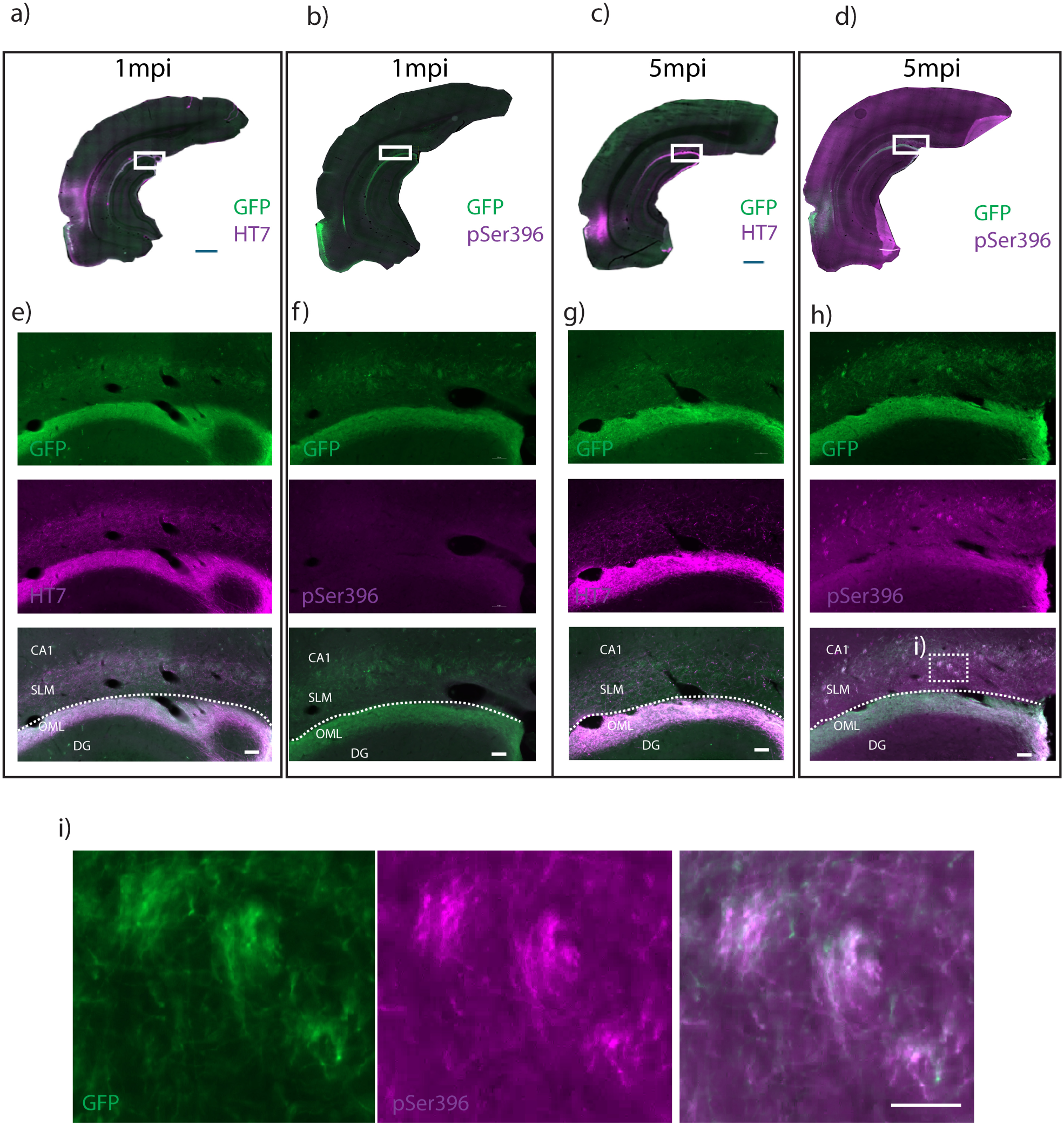
Following Layer II and III projections into the hippocampal formation reveals pSer396 labeling in projections belonging to LII and LIII neurons. Sections from 1MPI (left) and 5MPI (right) brains stained with GFP and HT7 (a,c,e,g) or pSer396 (b,d,f,h,i). **a.-d.)** Anatomical overview micrograph, Scalebar = 1000 um **e)** Micrograph from hippocampal formation (position indicated in a), note strong GFP label of the outer molecular layer of the dentate gyrus (OML) and the weaker GFP labeling in stratum lacunosum moleculare (SLM) of CA1, and near-identical label for HT7 (magenta). Hippocampal fissure is denoted by the dotted line. Scalebar= 50 um. **f)** Micrograph from 1 MPI labeled with pSer396 (magenta) showing no label at 1 MPI. Scalebar= 50 um **g)** Micrograph from 5 MPI labeled with HT7, note the HT7 signal both in OML DG and SLM CA1. Scalebar= 50 um. **h)** Micrograph from 5 MPI labeled with pSer396. Note the presence of pSer396 labeling in the OML of DG and the labeling in SLM CA1. Scalebar= 50 um. **i)** shows inset from **h)** highlighting the “Tuft” like pSer396 positive processes in CA1 SLM. Scalebar= 20 um. DG = dentate gyrus, CA1 = cornu ammonis 1.

#### hTau challenge leads to Reelin depletion in LEC LII at later stages

The final stage of AD-related tauopathy is of course neuronal death. While we did not see any signal with traditional markers of apoptosis (data not shown), there are reports that the neurodegeneration in AD does not actually proceed via apoptosis (Balusu and De Strooper 2024). Nevertheless, there are data consistent with significant cell death around the injection site. Neurons in the injection area appeared to have disrupted morphologies at 5 MPI (particularly in EC layers II and III) (**Supplemental figure 3-5**). At 1mpi the reelin positive layer appears the same between the injected and non injected sides (**Fig7a-c)**. In contrast, in 5 MPI animals both the fluorescent GFP and Reelin signal in LEC LII was greatly decreased compared with earlier timepoints, and with the uninjected side (**Fig7d-f**). The simplest explanation would be a decrease in the number of Reelin+ neurons near the rhinal fissure on the injected side. To quantitate this neuron loss, we counted reelin positive neurons per ROI in injected vs contralateral sides in adjacent sections from 4 animals at 5 MPI. We saw an approximately 44% decrease of reelin positive neurons in the injected sides of the animals (**Fig7g**), while at 1 MPI, injections into LEC do not appear to affect the expression of reelin (**Fig7a-c**). Taken together, these data strongly suggest that many of the hTau+ LEC LII fan cells are degenerating, as they do in early AD.

## Discussion

Like so many others, this study was inspired by Braak Stages (Braak et al. 2006), the progression of pathological tau species revealed by the detailed pathological staging of AD patient brains. Having seen the disparate roles played by entorhinal neuronal subtypes in memory function (Ohara et al. 2021; Witter et al. 2017; Kanter et al. 2017; Guardamagna et al. 2025; Vandrey et al. 2020), we, like others (Kawles et al. 2023) reasoned that one thing that could explain the anatomical stereotypy of Braak Stages in AD would be if the cellular microenvironment of different neuronal subtypes may be more or less conducive to the generation of tau pathology. We explored this idea by performing a “tau challenge”: a quantifiable viral overexpression of wildtype human 4R tau protein in a variety of cortical neurons in wildtype rat brain followed by immunological assessment of the rate of its conversion to pathological tau species, much as has been done before (Dujardin et al. 2014; Dujardin et al. 2018; Tetlow et al. 2023), but with special emphasis on the neuronal subtypes of the lateral entorhinal cortex. The results clearly showed that different neuronal subtypes have markedly different responses to the overexpression of human tau protein. Consistent with the pivotal role for the entorhinal cortex in the early development of AD-related tauopathy in prodromal patients, entorhinal neurons developed pathological tau species at a much faster rate than any other cortical neurons tested. However, even within the entorhinal cortex different neuronal subtypes had very different rates for the development of different pathological tau species. In particular, LEC LII fan cells) showed a strong predilection towards developing pathological tau species in response to hTau overexpression, with the vast majority of AT8+ neurons at 1 month post-injection being fan cells. An important caveat is that they also, however, tended to get greater overall expression levels as well. The finding that a subset of LEC LII and LEC LIII neurons showed an even greater proclivity to generate pathological tau species was far more surprising. These two neuronal subtypes also had markedly different tendencies to develop different kinds of pathological tau species, with LEC LII fan cells being more likely to develop AT8 positivity, while at least some LEC LIII neurons had a much higher per-unit hTau tendency to develop reactivity to the conformational epitope MC1 and later-stage phospho-antigen pSer396 relative to AT8. Moreover, we saw a population of perirhinal neurons expressing high AT8 levels around 2mpi. In sum, different neuronal subtypes clearly differ in their tendency to make pathological tau species from overexpression of wildtype human tau.

There are, of course, a number of limitations to these experiments. We are expressing hTau at far above physiological levels, which certainly helps explain how rapidly the pathology develops in our young animals, as overexpression of certain proteins can overwhelm chaperones and quality control processes (Sinnige, Yu, and Morimoto 2020). However, this is arguably the point of the experiment: by monitoring the level of hTau (via GFP) in every cell we can ask whether pathological tau species develop differentially (in rate and kind) in different neuronal subtypes per unit of hTau expression, and the answer, at least in wildtype rat brains, is clearly yes. Of course, the extent to which subtypes of cortical neurons are conserved between rodents and humans remains an ongoing question (Hodge et al. 2019), but the entorhinal cortex is one of the evolutionarily oldest cortical regions (periallocortex rather than neocortex (Insausti et al. 2017)) with many conserved structural and functional similarities between mammalian species (Witter et al. 2017). However, the most important caveat is that we are injecting wildtype human tau into young wildtype rat brains, and AD is a disease of aging involving normal levels of tau protein expression. The genetic linkage between the MAPT (i.e. tau) gene and AD is weak at best, so the only thing that connects our experiments to AD is the pivotal role of brain region of interest, the entorhinal cortex, in the earliest stages of AD.

Nevertheless, the fact that different neurons have markedly different innate tendencies to develop distinct pathological tau species could go a long way towards understanding its stereotypical progression of NFTs in AD and, indeed, tauopathies in general. A large variety of tauopathies exist, but most (e.g. fronto-temporal dementia) seem to begin in a particular pathology-specific population of neurons and then spread to synaptically-connected brain regions (McGeachan et al. 2025) and they also differ in the fine structure of their aggregated tau species (Shi et al. 2021). Both of these observations argue for a role of the cellular microenvironment of particular neuronal subtypes in tauopathies. Of course, this could happen in two ways: 1) the tauopathy could start in a particular neuronal subtype and spread to connected neurons transsynaptically (Gibbons, Lee, and Trojanowski 2019); or 2) different neurons could have different cell-autonomous intrinsic tendencies to develop pathological tau species, and the apparent transsynaptic transfer is largely coincidental. There are good arguments for both views, with tau seeding (Clavaguera et al. 2009; Clavaguera et al. 2013) being one of the most compelling arguments for the former. Support for the latter view comes largely from known anatomical inconsistencies, such as the fact that while the pre-alpha cells of Braak Stage I do project to the hippocampus, they project to the dentate gyrus and CA3, which are only implicated much later (Braak stage IV, V): Braak Stage II-III involves CA1 and subiculum, so this pathology cannot result from direct transfer from the pre-alpha neurons (Braak and Braak 1991).

Our results support both of these hypotheses, and remind us they are not mutually exclusive: different neuronal subtypes could have markedly different intrinsic tendencies towards the development of pathological tau species *and* the spread of the tauopathy could be largely transsynaptic. Indeed, even the tendency for the tauopathy to spread could be a subtype-specific property. This is consistent with our results showing that MC1 conformation, which has been suggested to be particularly pathogenic (Falcon et al. 2015; d’Abramo et al. 2013) and can even distinguish AD from (Falcon et al. 2015) Chronic traumatic encephalopathy (Sorrentino et al. 2023), is particularly evident in a subset of ECLIII neurons. If the human entorhinal cortex had a similarly hypersensitive ECLIII neuronal subtype it would explain why even though the disease starts in ECLII neurons (Braak Stage I) and then progresses to other neurons in the EC (Braak Stage II), the first AD-related tauopathy outside of the EC is not in CA3 and dentate gyrus, where ECLII neurons project in both rodents and humans, but CA1 and subiculum, where EC LIII neurons project (Witter et al. 1988). Different neuronal subtypes could thus be thought of as different biochemical contexts with a greater or lesser proclivity towards developing pathological tau species. There is precedent for this view: much of the recent success in oncology in recent years has owed to people studying not Cancer, but *cancers*… i.e. which pathway goes wrong in the context of which cell type (Pfohl et al. 2021). Recent developments in our understanding of the cellular complexity of the brain (Yao et al. 2021) and in the generation of vectors capable of distinguishing between neuronal subtypes (Blankvoort et al. 2018; Blankvoort, Descamps, and Kentros 2020; Nair et al. 2020; Hunker et al. 2025; Mich et al.) will not only lead to more accurate animal models of many disorders, it raises the exciting possibility of therapeutic interventions specifically targeting the particular neuronal subtypes that are most involved in a disorder, thereby minimizing negative side effects.

## Acknowledgements

The MC1 antibody was acquired through an agreement with Albert Einstein College of Medicine

Made available by the late P.Davies. We also want to thank Lluis Camprubi for suggesting and lending us a sample of this antibody.

We also thank Rajeevkumar Nair and the Viral Core Facility for help designing and producing viral vectors, and Qiangwei Zhang.

## Materials and Methods

### Animals

In the study we used Wistar rats. The animals were bred at the Kavli Institute for System Neuroscience at the Norwegian University of Science and Technology (NTNU). All experimental protocols were approved by the Norwegian Food Safety Authority (permit no. 30027) and are in accordance with the European Convention for the Protection of Vertebrate Animals used for Experimental and Other Scientific purposes.

### Molecular Cloning Methods

All molecular cloning methods including restriction digestion, ligation and DNA electrophoresis in agarose gel were performed according to the standard procedures. We generated two AAV constructs carrying two different isoforms of the human wild type Tau called respectively pAAV-hSyn-3R0N-GFP and pAAV-hSyn-4R0N-GFP. Each human Tau isoform is under the control of the human synapsin promoter (hSyn), and it is fused to the GFP as a reporter gene. The DNA insert expressing hSyn-4R0N or hSyn-3R0N have been synthetized from GenScript and cloned into the AAV backbone carrying the GFP cassette. The hTau-4R0N or the hTau-3R0N and the AAV backbone were separately digested with restriction enzymes provided by New England BioLab (NEB) and the ligation was performed according to the instructions provided by Rapid DNA Ligation Kit (Roche). After the ligation reaction, the DNA was used to transform the One Shot Stbl3 (Invitrogen) chemical competent bacteria using heat-shock procedures. Plasmid DNA extraction and purification was done by using QIAprep Spin Miniprep Kit (Qiagen) or HiSpeed Plasmid Kit (Qiagen). DNA fragments were isolated and purified from 1% agarose gel with QIAquick DNA gel extraction Kit (Qiagen).

### AAV production and Titering

Recombinant AAV vectors carrying the hΤau4R0N were packaged in AAV serotype 2/1 (<u>Hauck, Chen, and Xiao 2003</u>) and purified using the Heparin column affinity purification method (<u>McClure et al. 2011</u>). Briefly, the pAAV-containing the transgenes, pHelper, pRC (#240071, Agilent, USA) and pXR1 (NGVB, IU, USA) were transfected into Hek293T cell line (ATTC) using Lipofectamine transfection method (Thermo Fischer Scientific). Two days after the transfection the Hek293T cells containing the virus particles were scraped from the petri dishes, then isolated by centrifugation at 200Xg for 10 minutes. 150mM NaCl-20mM Tris pH8 buffer containing 10% sodium deoxycholate was added to the cell pellet for the lysis of the cells. The lysate was treated with benzonase nuclease HC (#71206-3, Millipore) at 37C for 45 minutes. Benzonase-treated lysate centrifuged at 3000 x g for 15 minutes, then subjected to HiTrap® Heparin High Performance (#17-0406-01, GE) affinity column chromatography using a peristaltic pump. The elute containing the viral particles was collected using Amicon Ultra centrifugal filters (#Z648043, Millipore). To calculate the number of genome-containg particles of the AAVs, we first made a standard curve using 102–108 copies of linearized plasmid. The purified AAV and standard curve DNA samples were quantified by qPCR using Power SYBR™ Green PCR Master Mix (#4368577, Thermo Fisher, USA) and WPRE (Fw WPRE 5’-CCGTTGTCAGGCAACGTG, Rw WPRE 5’-AGCTGACAGGTGGTGGCAAT) or ITRs (Fw ITR 5’-GGAACCCCTAGTAGTGGAGTT, Rw ITR 5’-CGGCCTCAGTGAGCGA) as primers.

### Stereotaxic Injection of AAV

The rAAVs were stereotaxtically injected into 4-5 weeks Wistar rats. 1 µl of rAAV was injected into the EC and S1 of the rats using the following coordinates: AP 2,29 Lambda, ML 6,5 DV 5 and AP 0,9 Bregma ML 3,8 DV 1,2 for S1. The rats were first anaesthetized with isoflurane gas and then the head was fixed to the stereotaxic frame with ear bars. A single injection of 1 µl virus was conducted with 20 pulses of 50 nl over 30 seconds using a nanoliter injector (Nanoliter 2010, World Precision Instruments, Sarasota, FL, USA), controlled by a micro syringe pump controller (Micro4 pump, World Precision Instruments). After the injections was delivered the glass pipette was left for 10 min before retracting.

### Perfusion

The rats were sacrificed 1 month, 2 months, and 5 months after rAAV injections. The animals were first anaesthetized with pentobarbital and then perfused transcardially with 0.9 % saline followed by 4 % paraformaldehyde in 0.1 M Phosphate buffer (pH 7.4) with 0.9 % saline. The brains were stored at 4C in 4% PFA overnight before being transferred to 1% DMSO solution containing 10% Glycerin in 0.125 M phosphate buffer for two or three days.

### Histology

Coronal rat brain sections of 30 µm were prepared using a sliding-freezing microtome at –30°C. Brain sections were stored at –20°C in 0.125 M phosphate buffer containing 40% Glycerin and 10% DMSO. Slightly different immunolabeling protocols were used for different Tau antibodies. For the immunostaining with pSer396 the sections were baked for antigen retrieval at 60°C for two hours and blocked for 1 hour at room temperature using PBS containing 0.1 % Triton X-100 and 10 % normal goat serum (PBT buffer). Sections were subsequently incubated with primary antibodies in PBT buffer overnight at 4°C with mild shaking. The day after, the PBS-washed sections were incubated for 2 hours at room temperature with secondary antibodies diluted in PBT buffer containing 5 % normal goat serum. For immunolabeling with all other tau antibodies, sections were heated at 60°C in PB buffer for 2 hours, and subsequently blocked in TBS 0.1% saponin with 10% normal goat serum. This was followed by washes in TBS saponin and subsequent antibody incubation for 18 hours with mild shaking. The following day, sections were rinsed and incubated with secondary antibodies for a minimum of 2 hours RT. After washing, sections are mounted with Diazabicylo[2.2.2] octane/polyvinyl alcohol (DABCO/PVA) onto Polysine slides (Menzel-Glaser, ThermoFisher, USA).

For 3,3’-Diaminobenzidine (DAB) staining, sections were heat-treated for 2 hours 60°C in PB buffer, quenched in 3% hydrogen peroxide (Merck Life Science AS/Sigma Aldrich Norway AS), blocked in TBS saponin with 10% normal goat serum and then incubated with primary antibody (AT8, 1:500) overnight. The following day, sections were rinsed in TBS saponin and incubated with secondary biotinylated antibody (1:1000). After rinsing, sections were incubated with the Avidin–Biotin complex (ABC, Vector Laboratories Cat# PK-4000) for 90 minutes RT. After final rinses, sections were stained with the DAB chromogen and after rinsing mounted on Polysine slides left to dry overnight and sealed with Xylene/Entellan mix.

### Imaging and quantification

All tissue sections were scanned on an automated Zeiss Axio Scan.Z1 system with a 20 × objective (Plan-Apochromat 20x/0.8 M27). Analyses and quantifications were done in QuPath(Bankhead et al. 2017). Briefly, images were loaded into a project, the brush tool was used to define ROIs. Most often injections involvedLEC LII, LIII, LEC LIV-VI, BA35 and BA36, though in some animals the injection included piriform cortex LII and LIII or CA1 as well, and these were always included in the ROI.We used the cell detection feature, using GFP (thus all objects detected are positive for GFP) (or Reln for Fig7.) as the signal to define cell bodies. Detection measurements were then exported to excel and analyzed using custom Jupyter notebook scripts and GraphPad Prism. Briefly, Fluorescence in mean fluorescence in AF488 (GFP) and mean fluorescence in AF647 (Tau antibodies) are plotted against each other. The custom Jupyter script loads and normalizes data per animal, antibody and timepoint. Linear regressions were computed and a regression line was plotted for each timepoint and antibody. Slopes derived from individual animals were statistically compared to a theoretical 1:1 relationship between Tau and GFP, using a two-sided one-sample t-test. For classification a hypothetical X=Y was imagined with a 2SD interval around it. Objects outside where classified as “Above” or “Below”, the “Above” class was further subdivided into High-Tau (thresholded at a normalized Tau signal of 5 and above). The Objects within 2SD of the linear relationship were subdivided and labeled as Linear-Low Tau (hTau<3), Linear Mid-tau(hTau3-5) and Linear High-Tau (hTau > 5). Then classes were checked for the contribution of different anatomical areas (Parents) to the composition of classified objects.

For Reelin quantification, injected and non-injected sections of comparable cortical AP position were analyzed. Regions of interest (ROIs) covering the dorsal aspect of LEC LII were manually segmented on randomized images with the GFP channel hidden. Cells labeled by Reelin (AF647 channel) were detected using the QuPath Cell Detection tool. The mean number of Reelin-positive neurons on the injected side was compared to the mean number on the contralateral, un-injected side for each animal. Students paired t-test was used to analyze the difference between both sides. GraphPad Prism was used to analyze the Reln+ neuron counts and generate the graphs for Fig 7. Students paired t-test was used to analyze the difference between both sides. GraphPad Prism was used to analyze the Reln+ neuron counts and generate the graphs for Fig 7.

**Figure 7.**
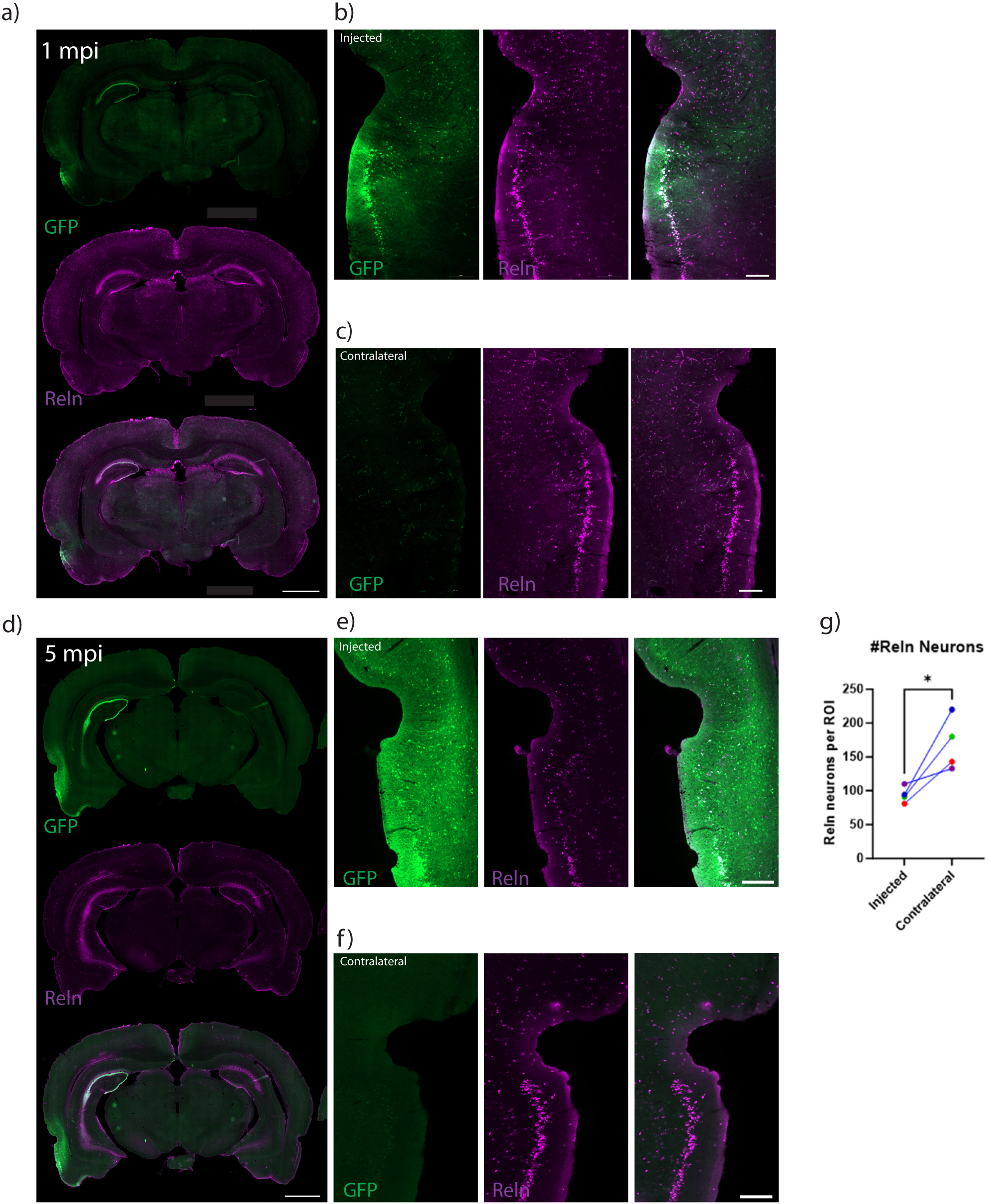
Reelin depletion in LII at 5MPI suggests hTau challenge leads to degeneration of LECLII fan cells. **a)** Representative whole brain section from 1 MPI labeled with reelin (Magenta) Scalebar 2000=um. **b)** inset from injected side rhinal fissure. Scalebar = 200um **c)** inset from contralateral rhinal fissure. Note that reelin neurons in layer II are similar between the two sides. Scalebar = 200um **d)** Whole brain section from 5 MPI animal. Scalebar 2000=um. **e)** Inset from injected side. Note the marked depletion of reelin neurons in LEC LII. Scalebar = 200um. **f)** Inset from contralateral side rhinal fissure. Note that reelin neurons are preserved on this non-injected side. Scalebar = 200um. **g)** Quantification of reelin positive neurons in LEC LII ROIs from injected vs contralateral sides. N = 4 animals. Mean injected side = 94/ROI, mean contralateral = 169/ROI. p = 0.0409 student’s paired T-test.

### Antibodies

The primary antibody used were rabbit anti 4-R-Tau [clone EPR21725](Abcam, 1:1000), rabbit anti-phospho Tau396 (Clone EPR2731, Abcam, 1:2000), rabbit anti-phospho TauAT8 (Nordic Biosite, 1:500), chicken anti-GFP (Abcam, 1:1000), mouse anti human Tau(HT7, Fischer Scientific, 1:1000) and mouse anti-Reelin (Millipore, 1:1000), rabbit anti-WFS1(1:1000) mouse anti Calbindin D28K (1:1000).

All corresponding secondary antibodies were from ThermoFisher/Life technologies or Jackson Immuno Research laboratories, USA, used at a dilution of 1:1000.

## Notes

### Competing Interest Statement

The authors have declared no competing interest.

### Funding

- The Research Council of Norway: 332640, 197467, 249945
- Trond Mohn stiftelse
- Nasjonalforeningen for Folkehelsen: 35278

