## Supplemental figure+legends} for "Preferential generation of pathological tau species in specific subtypes of entorhinal neurons: implications for Alzheimer’s Disease"

Supplemental figure 1.

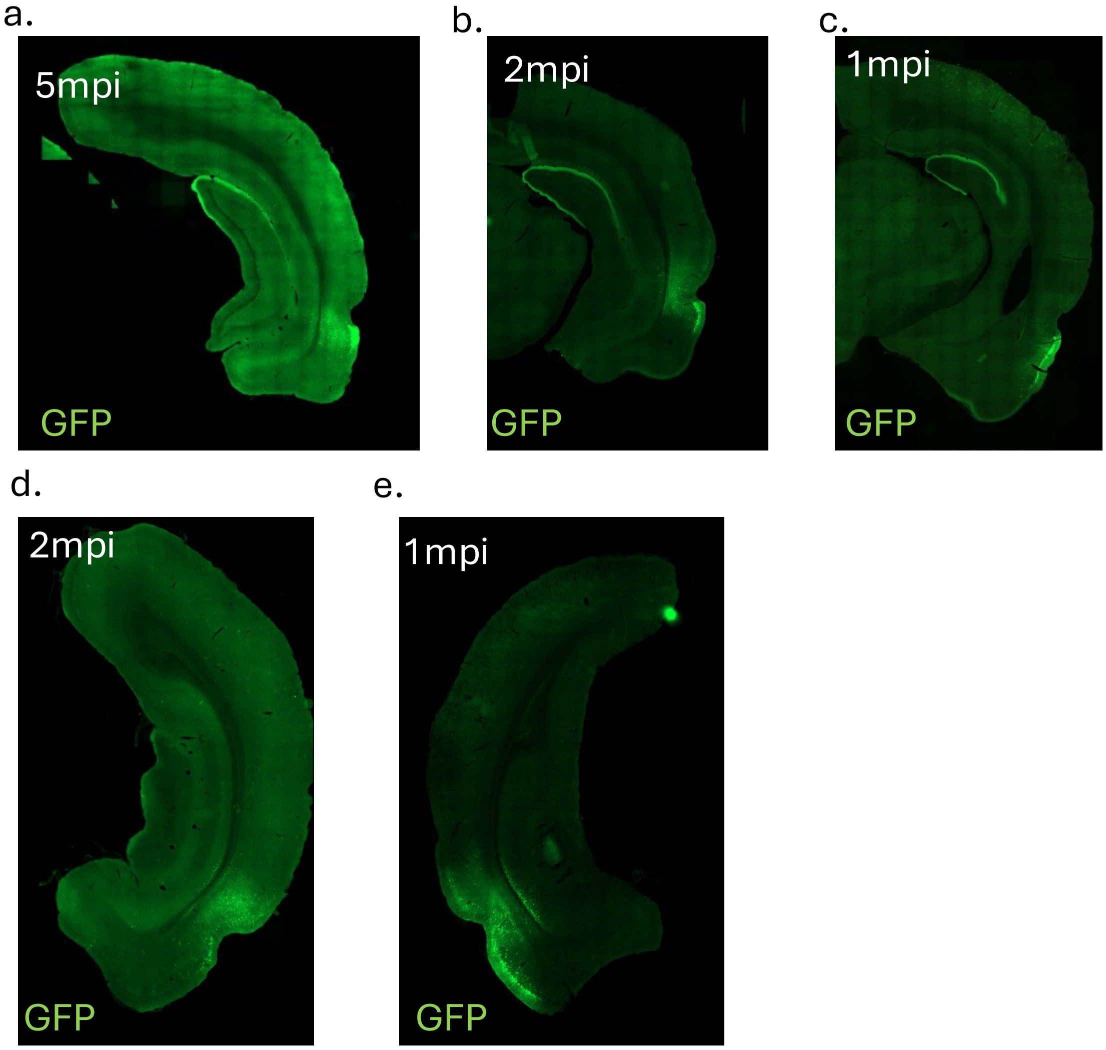


**Supplemental figure 1. Example injections GFP across rhinal cortices. A)** Micrograph from a 5mpi animal. Nothe that GFP is present both above and below the rhinal fissure albeit at varying degrees. **B)** micrograph from 2mpi animal, Injection around rhinal fissure with substantial signal also in cortex above the rhinal fissure. **C)** micrograph from 1mpi animal where the injection mainly targets LEC and Piriform cortex. **D)** micrograph from 2mpi animal with ectorhinal centric GFP expression. **E)** micrograph from 1 mpi animal with injection area stretching form association cortices to ventral LEC.

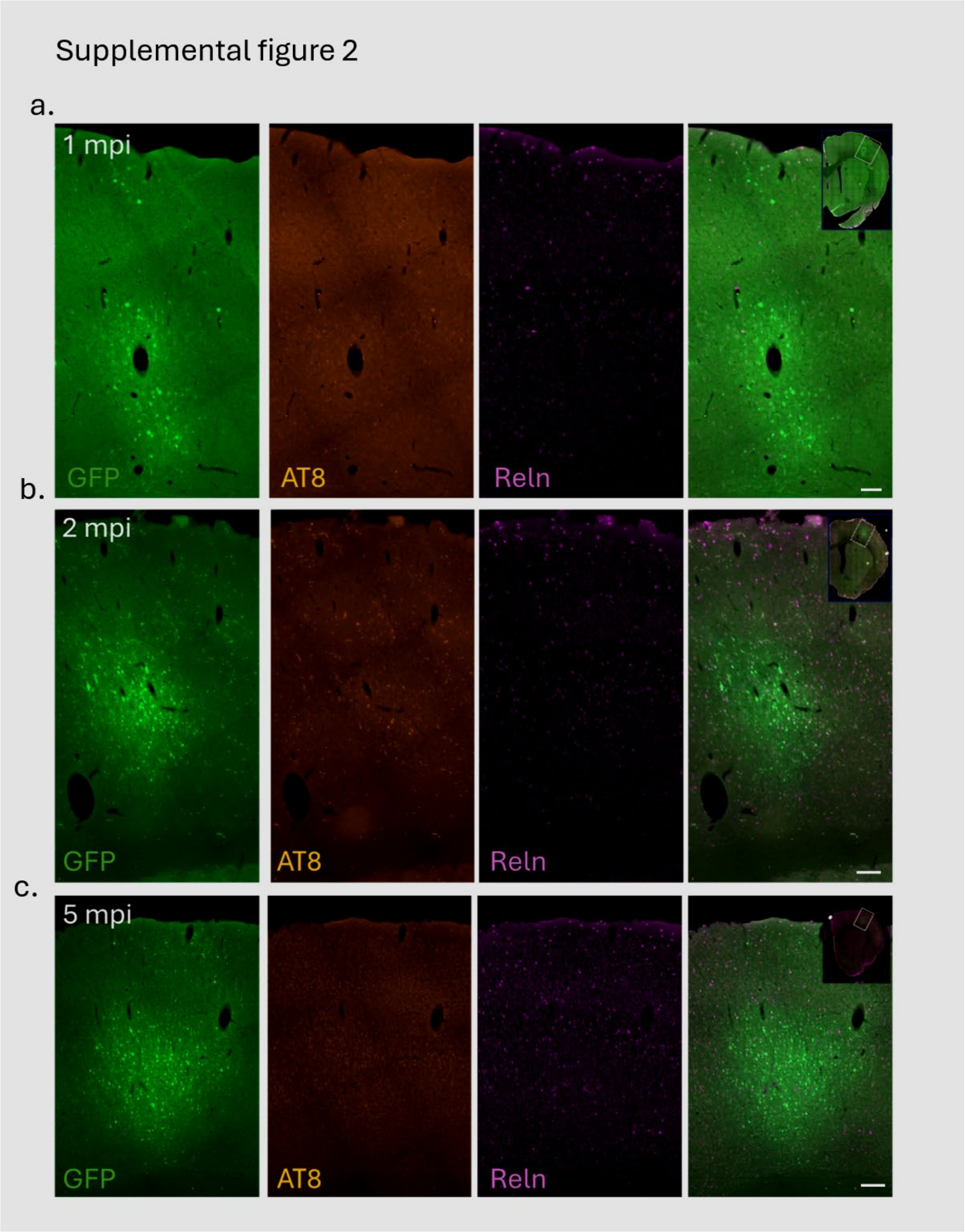


**Supplemental figure 2. S1 injections do not show substantial pTau expression. A)** micrograph of S1 injection site 1mpi. Labels are GFP (green), AT8 (Orange) and Reelin (Magenta). Upper right inset shows the whole section localization. Scalebar = 100um. **B)** as **A)** but 2 mpi injected animal. **C)** 5 mpi S1 injection. Note how AT8 labeling is relatively absent and mostly driven by background fluorescence.

Supplemental figure 3

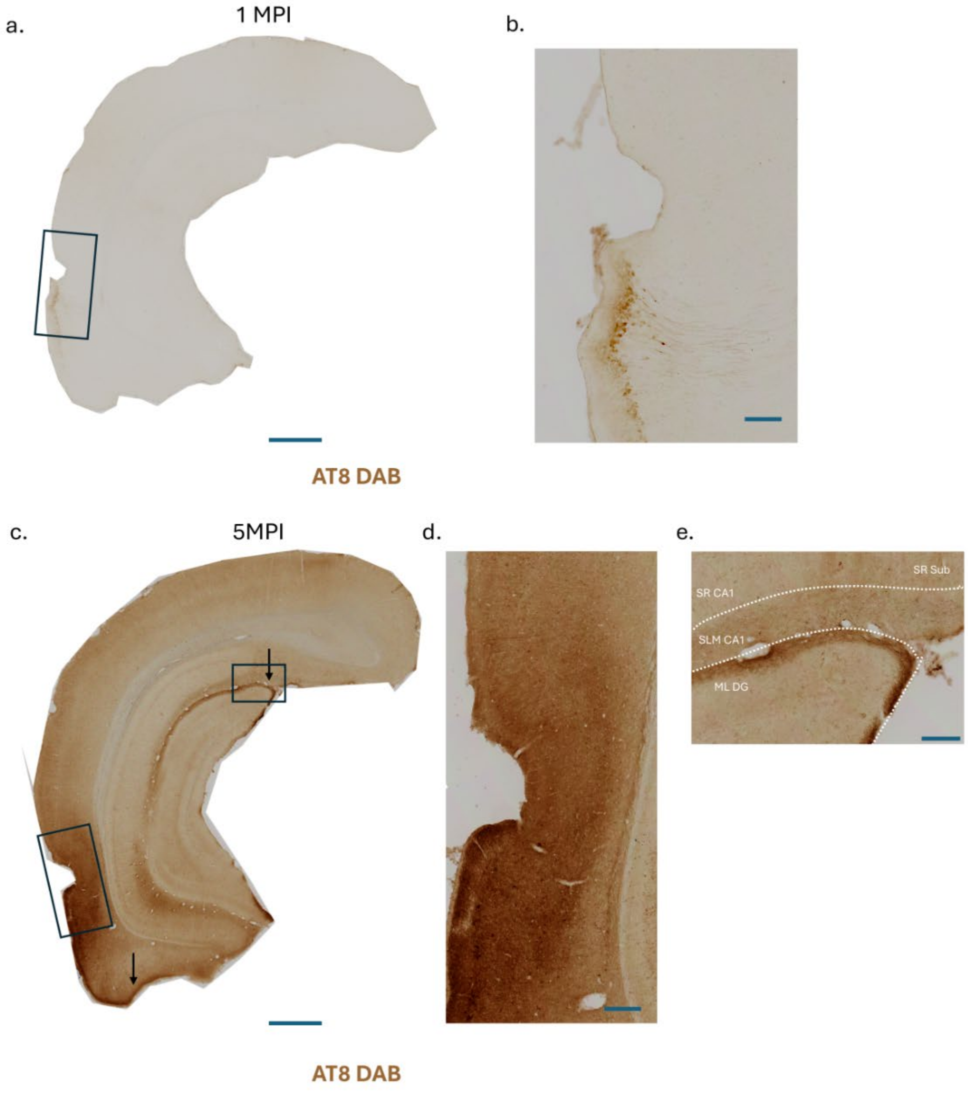


**Supplemental figure 3. DAB labeled 1mpi and 5mpi animals reveals progressive AT8 Tau pathology and spread from LII to hippocampal neurites.** **A)** Micrograph of 1mpi injected animal stained with AT8 DAB. Note that AT8 signal at this timepoint is largely localized to LEC LII. Scalebar = 1000um **B)** inset from **A)** showing strong AT8 signal in LEC. Scalebar = 200um. **C)** AT8 labeled 5 mpi animal highlight how AT8 signal increases around the rhinal fissure in a progressive manner. Arrows point to AT8 signal in hippocampal processes and LI projections to MEC. **D)** Inset around rhinal fissure shows presence of AT8 pTau in numerous cells and processes. Scalebar = 200um. **E)** inset from hippocampal region. The outer Molecular Layer (ML) of the dentate gyrus displays strong AT8 positive staining at this timepoint and originates from LII Fan-cells. Note AT8 signal in the Stratum Lacunosum Moleculare (SLM) of CA1, this AT8 has a non-Fan-cell origin. Scalebar = 200um. Abb. SR = Stratum Radiatum. Sub = Subiculum.

Supplemental figure 4

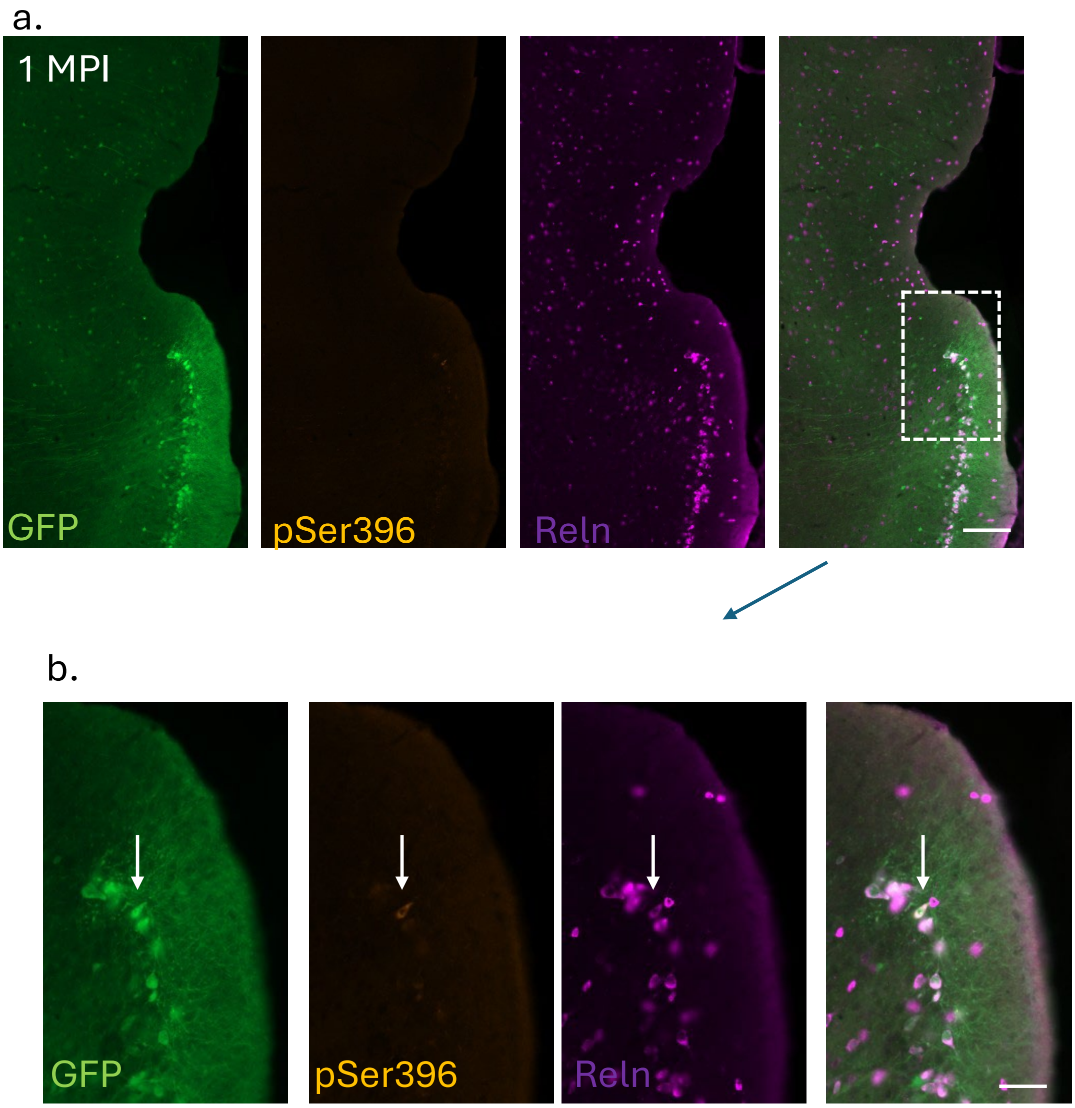


**Supplemental figure 4. Sparse pSer396 labeling at 1mpi LEC LII. A)** micrograph from 1mpi animal labeled with pSer396 and Reln. Scalebar 200um. **B)** inset from A) shows pSer396 positive neuron in LII of LEC. Note that it is a single Reln positive neurons with similar GFP expression to its neighbors. Scalebar = 100um.

Supplemental figure 5

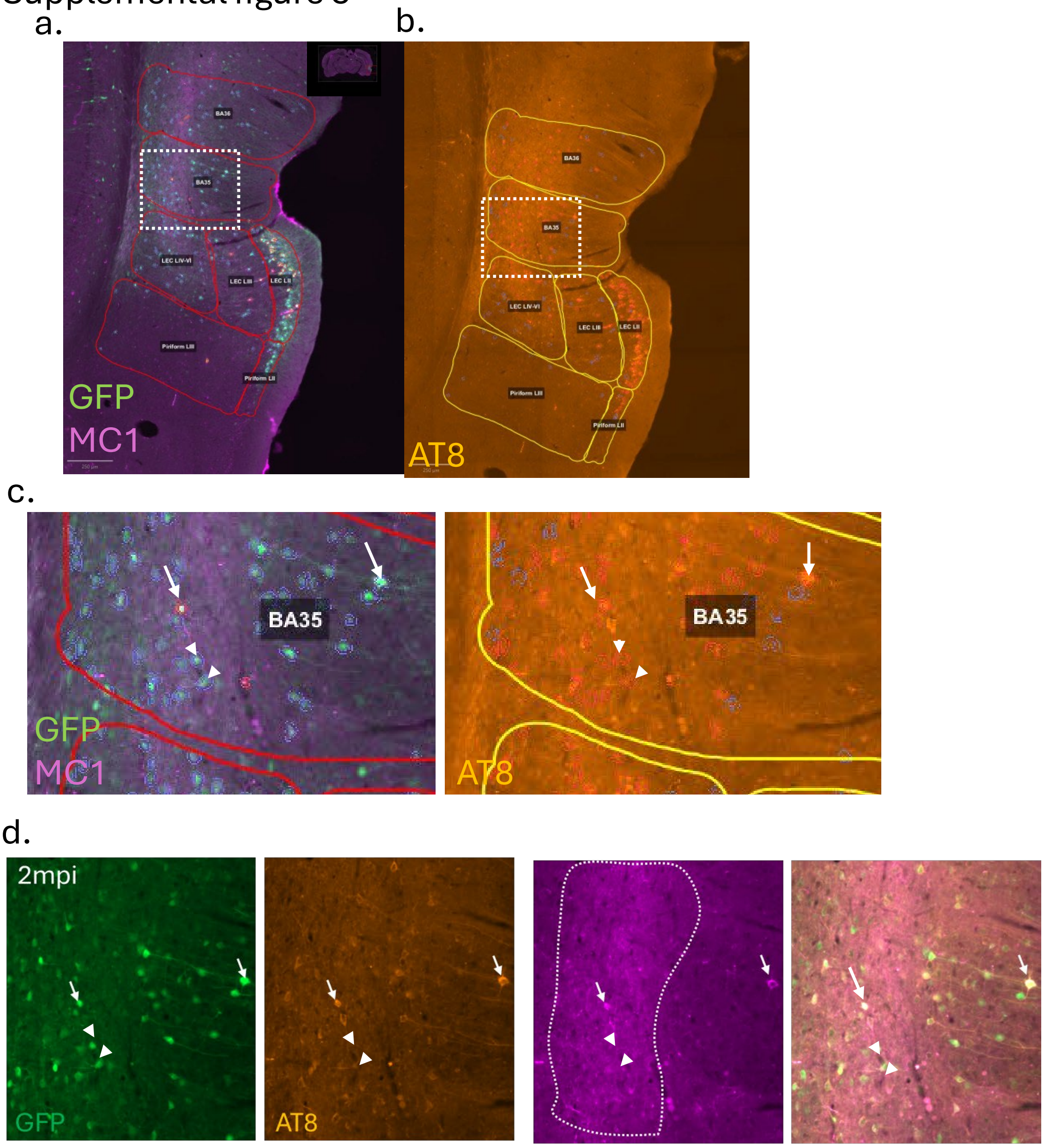


**Supplemental figure 5. High tau signal in tau neurites in BA35 at 2 mpi drives hTau fluorescence in deeper layers. A)** QuPath output from GFP positive neurons detected around the rhinal fissure. Parent regions are labeled and detected cell objects are circles. Red ROI masks indicate neurons with high MC1 signal whereas blue ROI mask indicates neurons below threshold MC1 signal. Scalebar = 250um. GFP (GFP) and MC1 (Magenta). **B)** QuPath output for GFP detected cell masks in the same area, but labeled for AT8. Red ROI masks indicate high threshold AT8+ neurons and blue ROI masks indicate below threshold objects. **C)** insets from **A**) and **B)** showing detected neurons in BA35. Arrows show the same two neurons in both figures, these two neurons are both positive for AT8 and MC1. Arrowheads indicate neurons that due to high signal in neurites surrounding deeper BA35 are labeled as positive from QuPath although they do not have AT8 or MC1 in the cellbody. **D)** Magnification and higher quality image of same cells from **C),** Note that while many neurons in deeper BA35 are positive for AT8, MC1 is mainly seen in neurites, although the ROI masks for neurons will incorporate this “neuritic tau” into the total cell signal. The encircled area highlights the neurite tau signal that might inflate average fluorescence in deeper BA35 and some BA36 neurons.

### Supplemental figure 6

a.

b.

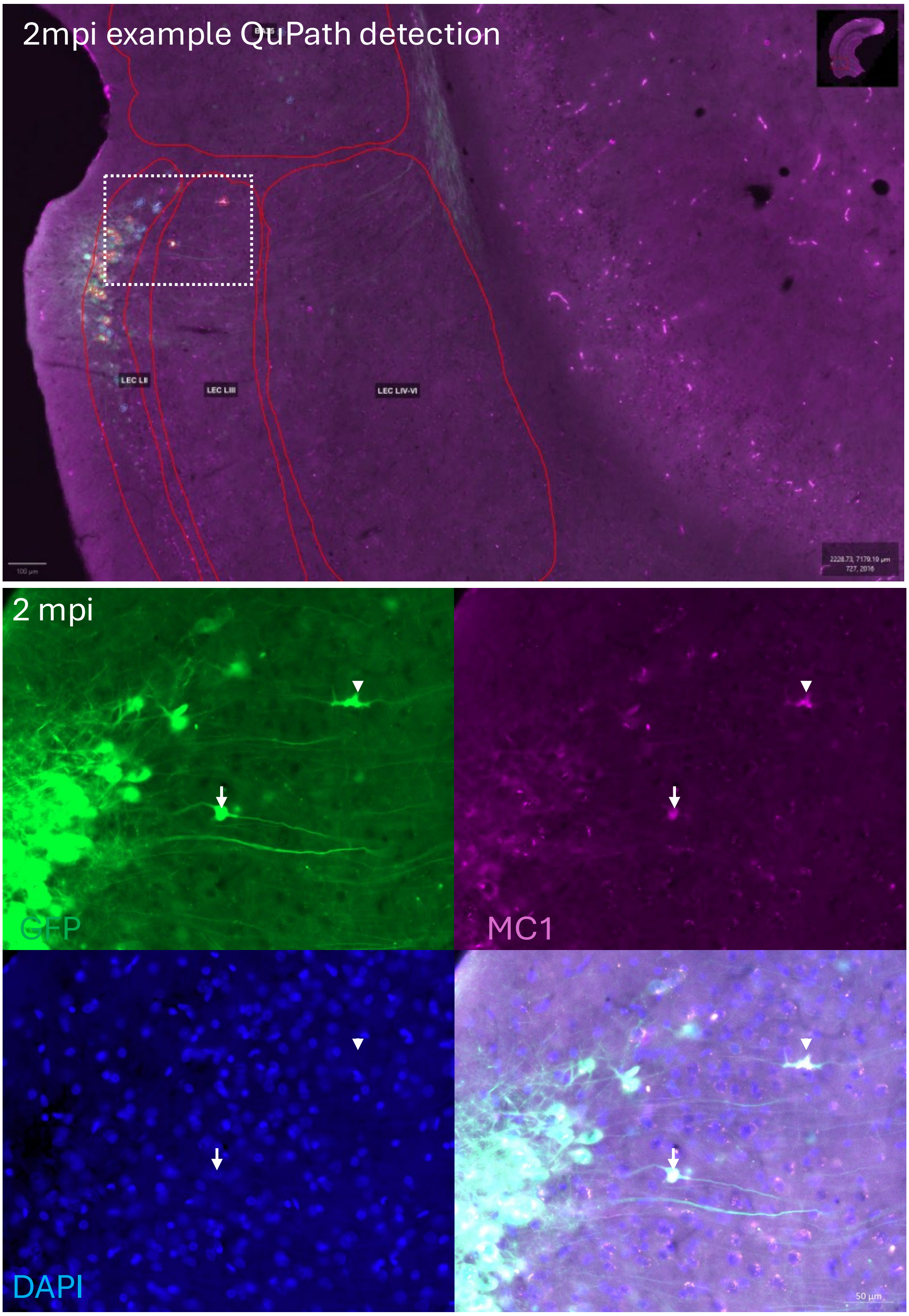

**Supplemental figure 6. Example of putative LIII neuron and dystrophic neurite. A)** Example output detections from QuPath. Micrograph is a merge of GFP(Green) and MC1(Magenta). Note ROIs for LEC LII, LIII, LIV-VI, BA35 and BA36. Circles around neurons highlight GFP detected neurons (Blue = below threshold Tau signal, red = Above threshold Tau signal). Scalebar= 100um. **B)** Higher magnification Inset of LII/LIII region. Arrowhead indicates putative axonal varicosity misidentified as neuron due to no DAPI signal. Whereas the arrow indicates putative neuron with high MC1 Tau in LIII. Scalebar = 50um.

Supplemental figure 7

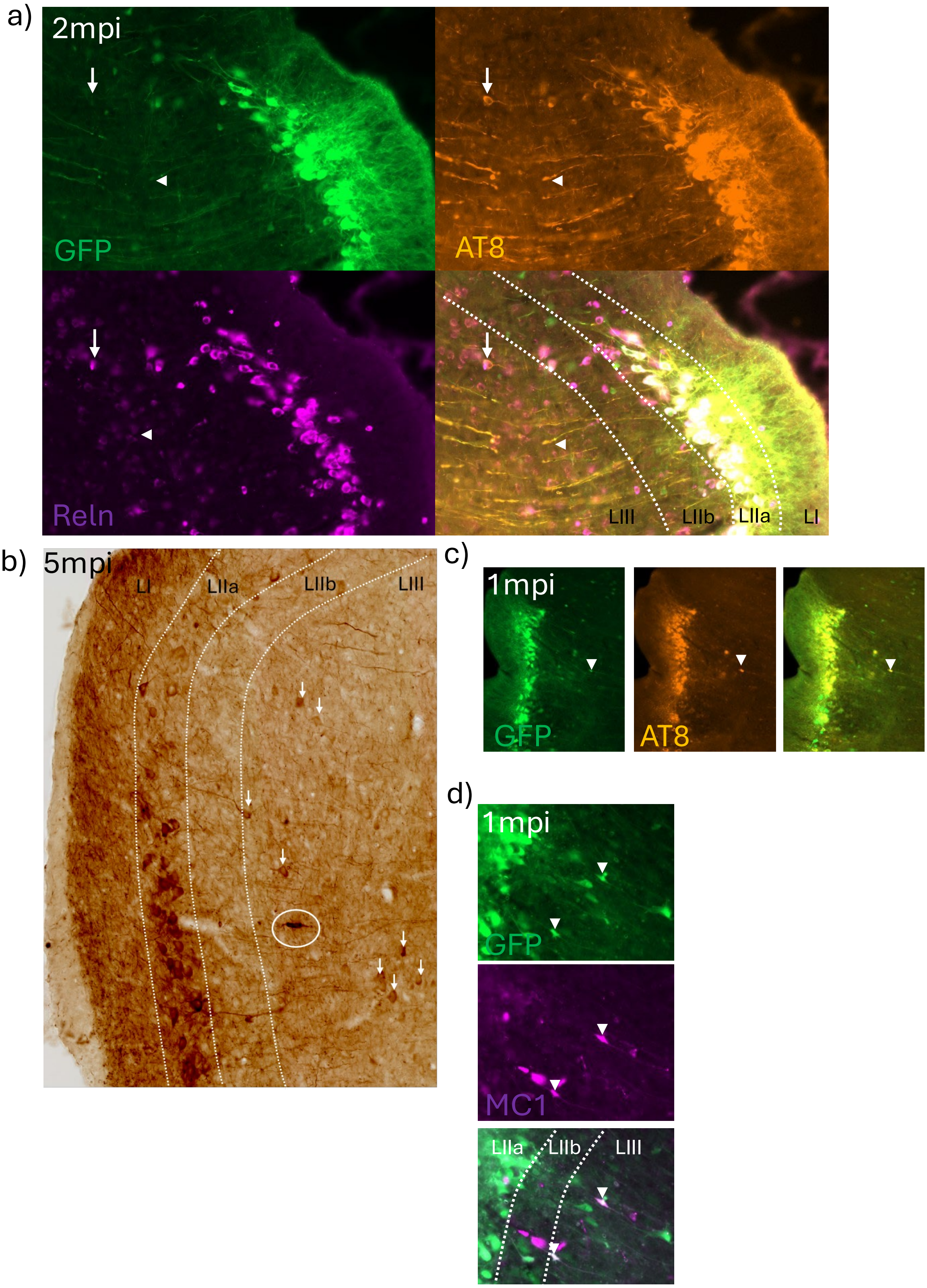


**Supplemental figure 7. Axonal varicosities and proper neurons in LIII. A)** a) micrograph from LEC LII/III at 2 mpi. Arrow highlight proper neurons positive for AT8 in LIII whereas arrowhead indicates

AT8 neurites that potentially could be misidentified as neurons. **B)** Micrograph from 5 mpi animal labeled with AT8 DAB shows presence of proper AT8 positive neurons in LIII highlight by arrows, circled is a potential axonal varicosity/dystrophy which could be misidentified as a neuron. **C)** micrograph from 1 mpi animal, arrowhead shows potential artefact in LIII. **D)** micrograph from 1 mpi animal shows MC1 positive neurite swellings in LIIb/LIII.
